# Antibody Escape by Polyomavirus Capsid Mutation Facilitates Neurovirulence

**DOI:** 10.1101/2020.07.14.203281

**Authors:** Matthew D. Lauver, Daniel J. Goetschius, Colleen S. Netherby-Winslow, Katelyn N. Ayers, Ge Jin, Daniel G. Haas, Elizabeth L. Frost, Sung Hyun Cho, Carol M. Bator, Stephanie M. Bywaters, Neil D. Christensen, Susan L. Hafenstein, Aron E. Lukacher

**Affiliations:** Department of Microbiology and Immunology, Penn State College of Medicine, Hershey, PA 17033, USA; Department of Biochemistry and Molecular Biology, Pennsylvania State University, University Park, PA 16802, USA; Huck Institutes of the Life Sciences, Pennsylvania State University, University Park, PA 16802, USA; Department of Pathology, Penn State College of Medicine, Hershey, PA 17033, USA; The Jake Gittlen Laboratories for Cancer Research, Penn State College of Medicine, Hershey, PA 17033, USA; Department of Medicine, Penn State College of Medicine, Hershey, PA 17033, USA

**Keywords:** Polyomavirus, Neutralizing antibody, Progressive multifocal leukoencephalopathy (PML), Cryo EM, Conformational epitope, Subvolume refinement, Fab:Capsid complex

## Abstract

JCPyV polyomavirus, a member of the human virome, causes Progressive Multifocal Leukoencephalopathy (PML), an oft-fatal demyelinating brain disease in individuals receiving immunomodulatory therapies. Mutations in the major viral capsid protein, VP1, are common in JCPyV from PML patients (JCPyV-PML) but whether they confer neurovirulence or escape from virus-neutralizing antibody (nAb) *in vivo* is unknown. A mouse polyomavirus (MuPyV) with a sequence-equivalent JCPyV-PML VP1 mutation replicated poorly in the kidney, a major reservoir for JCPyV persistence, but retained the CNS infectivity, cell tropism, and neuropathology of the parental virus. This mutation rendered MuPyV resistant to a monoclonal Ab (mAb), whose specificity overlapped the endogenous anti-VP1 response. Using cryo EM and a custom subvolume refinement approach, we resolved an MuPyV:Fab complex map to 3.1 Å resolution. The structure revealed the mechanism of mAb evasion. Our findings demonstrate convergence between nAb evasion and CNS neurovirulence *in vivo* by a frequent JCPyV-PML VP1 mutation.

## INTRODUCTION

The humoral immune response is critical for controlling acute and persistent viral infections; evasion of the neutralizing antibody (nAb) response often underlies virus-mediated morbidity and mortality. Seasonal influenza vaccinations are necessitated by the emergence of influenza A virus subtypes with mutations in hemagglutinin and neuraminidase capsid proteins that handicap neutralization by virus-specific antibodies (Bedford et al., 2015; Hensley et al., 2009; Petrova and Russell, 2018). Viruses causing persistent infections also acquire mutations that evade nAbs (Ciurea et al., 2000; Inuzuka et al., 2018; Kinchen et al., 2018; Salpini et al., 2015; Wei et al., 2003). The human virome is comprised of a sizeable number of persistent viruses whose pathogenicity is restrained by a healthy adaptive immune system (Virgin et al., 2009).

JC polyomavirus (JCPyV) is a prevalent member of the human virome (Kamminga et al., 2018; Viscidi et al., 2011). Immunological perturbations are a necessary antecedent for progressive multifocal leukoencephalopathy (PML), a fatal demyelinating brain disease caused by JCPyV (Haley and Atwood, 2017). JCPyV isolates from the brain, cerebrospinal fluid (CSF), and blood of PML patients contain unique mutations not found in virus present in patients’ urine or in circulating (archetype) strains (Van Loy et al., 2015; Vaz et al., 2000). These JCPyV-PML variants contain rearrangements in the noncoding control region including deletions, insertions, and duplications. These rearrangements alter transcription factor binding sites and enhance viral replication in glial cells (Gosert et al., 2010; Marshall et al., 2010). In addition, most JCPyV-PML variants have non-synonymous mutations in VP1, the major viral capsid protein, with the most common being a leucine-to-phenylalanine substitution at residue 54 (L54F) and a serine-to-phenylalanine/tyrosine substitution at residue 268 (S268F/Y) (Gorelik et al., 2011). These VP1 mutations have been reported to alter viral receptor binding, resulting in the utilization of a restricted set of receptors for cellular attachment and entry, thereby altering viral tropism (Geoghegan et al., 2017; Maginnis et al., 2013). In hypomyelinated RAG^-/-^ mice engrafted with human glial precursor cells (GPCs), however, infection with wild type or VP1 mutant JCPyVs resulted in similar levels of glial cell infection (Kondo et al., 2014). Recent evidence has also implicated VP1 mutations as nAb escape variants. PML patient sera only weakly neutralized patient-matched JCPyV VP1 variants (Ray et al., 2015). nAbs recognize antigenic epitopes that may overlap with receptor-binding sites; therefore, capsid mutations can affect both cellular tropism and humoral immunity (Kinchen et al., 2018; Lynch et al., 2015; McKnight et al., 1995; Reh et al., 2018). The relative impact of VP1 mutations in JCPyV on nAb recognition and tissue tropism is unknown.

The tight species-specificity of polyomaviruses obviates investigating the role of these JCPyV-PML VP1 mutations *in vivo*. Mouse polyomavirus (MuPyV) shares many features with JCPyV, including asymptomatic persistent infection, viral persistence in the kidney, and control by the virus-specific adaptive immune response (Berger et al., 2017; Du Pasquier et al., 2004; Han Lee et al., 2006; Szomolanyi-Tsuda and Welsh, 1996). MuPyV and JCPyV are non-enveloped, circular dsDNA viruses with capsids ∼45 nm in diameter. Their 5 kb genomes encode the nonstructural T antigen genes and the VP1 and VP2/VP3 structural proteins. Five copies of VP1 intertwine to form each capsomer subunit that also incorporates one copy of VP2/VP3 (Hurdiss et al., 2016). The core secondary structure formed by VP1 is the antiparallel β barrel structure commonly called the jellyroll. The β strands (BIDG and CHEF) are connected by flexible loops (BC, DE, EF, and HI) that extend outward from the surface of the capsomer and comprise the majority of the VP1 hypervariable regions. VP1 C-terminal extensions interact to link capsomers together forming the T=7d icosahedron. Twelve capsomers lay on an icosahedral five-fold vertex surrounded by five neighboring capsomers and are referred to as pentavalent capsomers. Each of the remaining sixty capsomers in the icosahedron has six neighboring subunits and is referred to as a hexavalent capsomer (Caspar and Klug, 1962; Harrison, 2017; Hurdiss et al., 2018). Each asymmetric unit contains six VP1 molecules that are structurally distinct because they experience different environments. The five VP1 molecules within a pentavalent capsomer are structurally identical, whereas the five VP1 molecules within each hexavalent capsomer are quasi-equivalent (Caspar and Klug, 1962).

The X-ray structure of MuPyV VP1 pentamers has been solved at resolutions ranging from 1.64 Å to 2.0 Å (Buch et al., 2015; Stehle and Harrison, 1997). However, crystallization of isolated pentamers may not represent the native environment of the icosahedral capsid. There are several structures of the entire icosahedral capsid including a 3.65 Å resolution X-ray map for MuPyV and a 3.4 Å resolution cryo EM structure of BKPyV (Hurdiss et al., 2018; Stehle and Harrison, 1996). Notably a 4.2 Å map of BKPyV interacting with single chain variable fragment (scFv) provided insight into antibody neutralization of polyomavirus (Lindner et al., 2019). Structural studies typically use the fragment antigen-binding (Fab) to avoid potential cross-linking of capsids by and multiple points of intrinsic flexibility of intact antibodies that interfere with cryo EM analysis. Use of the Fab domain (or scFv) maximizes attainable resolution at the experimentally relevant antigen-binding domain.

Recent hardware advances in cryo EM have led to major innovations in software design that overcomes limitations resulting from particle flexibility, heterogeneity, and imperfect symmetry (Ilca et al., 2015; McMullan et al., 2016; Scheres et al., 2009). Subparticle refinement approaches have made possible higher resolution maps of large, flexible virus capsids (Bhella, 2019; Chen et al., 2018; Zhu et al., 2018). Atomic resolution structures of virus-Fab complexes can elucidate mechanisms of neutralization and define conformational epitopes on the capsid, including key residues involved in recognition by the antibody (Dong et al., 2017; He et al., 2020; Zhu et al., 2018). Virus-Fab complex structures may predict viable escape mutations that naturally emerge under selective pressure from nAbs.

S268 of JCPyV corresponds to V296 of MuPyV VP1, with the residues mapping to the same position on the capsid (Sunyaev et al., 2009). We found that MuPyV carrying the V296F VP1 mutation, was impaired in its ability to replicate in the kidney, but replicated in the brain equivalently to parental virus. In addition, this mutant virus was completely resistant to neutralization by a VP1 mAb (clone 8A7H5), which recognizes a VP1 region overlapping the dominant target of the endogenous antibody response (Swimm et al., 2010). To determine the mechanism of humoral escape we solved the cryo EM structures of MuPyV in the presence and absence of the fragment antigen-binding (Fab) of the VP1 mAb 8A7H5. Using a novel and rapid local refinement strategy, we attained atomic resolution and could build the Fab variable domain *de novo*. This cryo EM analysis identified unambiguous contact residues at the interface between VP1 and Fab, and revealed the mechanism of immune escape by the V296F variant. This atomic resolution description of the 8A7H5 epitope also provided plausible neutralization escape mechanisms for other JCPyV-PML mutations and several additional MuPyV variants isolated *in vitro*. Together, our data demonstrate that VP1 mutations in polyomaviruses concomitantly enabled evasion of the nAb response and facilitated neurovirulence by preserving viral replication in the CNS. These findings support the concept that viremia by nAb-resistant VP1 JCPyV variants is a critical early step in PML pathogenesis.

## RESULTS

### V296F VP1 mutation retains MuPyV tropism for brain but not kidney

To model the effects of the S268F PML mutation, we generated a MuPyV mutant containing V296F substitution in the wild type (WT) A2 strain (A2.V296F) (**Figure 1A**) (Dawe et al., 1987; Sunyaev et al., 2009). A2.V296F exhibited a slight reduction compared to WT virus in a 60 h single-cycle replication assay, but showed equivalent expression of the nonstructural Large T antigen (LT) mRNA 24 hours post-infection (hpi) in several mouse cell lines and primary mouse embryonic fibroblasts (**Figures S1A and S1B**). Thus, this VP1 mutation had little impact on the ability of MuPyV to infect and replicate *in vitro*.

**Figure 1.**
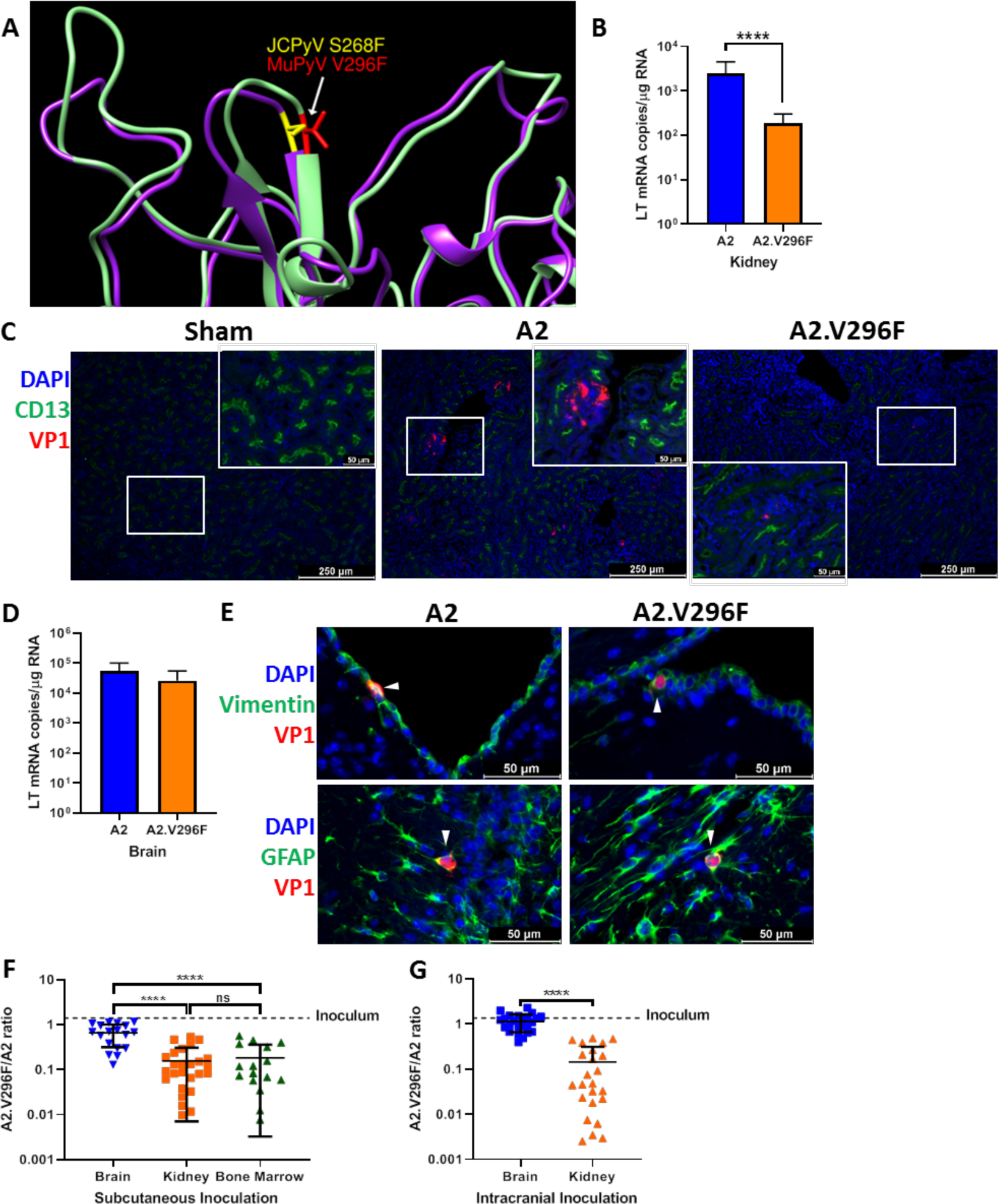
The V296F VP1 mutation in MuPyV impairs kidney, but not brain, infection. (A) Structural comparison of JCPyV.S268 (PDB 3NXG) and MuPyV.V296 (PDB 5CPU) VP1 residues (Buch et al., 2015; Neu et al., 2010). (B) A2 and A2.V296F LT mRNA levels 4 dpi in the kidneys of mice infected s.c. Data are from three independent experiments, n = 17 mice. (C) 100x images of kidney cortices from CD8-depleted, STAT1^-/-^ mice 7 dpi. Inset is a 400x image of the region outlined in white. Representative of two independent experiments. (D) A2 and A2.V296F LT mRNA levels 4 dpi in the brains of mice infected i.c. Data are from three independent experiments, n = 12-13 mice. (E) 400x images of brains 4 dpi with A2 or A2.V296F i.c. VP1^+^ cells are indicated with white arrows. Representative of three independent experiments. (F & G) Ratio of A2.V296F to A2 in various organs of mice 14 dpi with a 1:1 PFU inoculum of A2:A2.V296F s.c. (F) or i.c. (G). The dotted line indicates the ratio of A2:A2.V296F DNA in the inoculum. Data are from 2-3 independent experiments, n = 16-26 mice. Data were analyzed by Mann-Whitney *U* test (B, D, G) or one-way ANOVA (F). ****p<0.0001.

Little is known how the JCPyV S268F mutation affects JCPyV tropism *in vivo*. In PML patients, VP1 mutant viruses are detected in blood, CSF, and brain tissue, but not urine (Gorelik et al., 2011; Reid et al., 2011). Because the kidney is a reservoir for both JCPyV and MuPyV persistence, the absence of JCPyV VP1 mutant virus in the urine led us to ask whether the S268F virus exhibited a defect in kidney tropism. Compared to mice inoculated subcutaneously (s.c.) with parental A2, mice given A2.V296F showed significantly lower infection levels in the kidney 4 days post-infection (dpi) (**Figure 1B**). Immunocompetent mice infected with MuPyV do not develop productive kidney infection when detected by immunofluorescence or immunohistochemistry (Drake and Lukacher, 1998). STAT1^-/-^ mice depleted of CD8^+^ T cells develop severe systemic MuPyV infection (Mockus et al., 2020). We compared A2 and A2.V296F kidney infection in CD8^+^ T cell-depleted STAT1^-/-^ mice by staining kidney sections at day 7 pi for VP1 and CD13, which marks proximal tubules in the kidney cortex (Guder and Ross, 1984; Van der Hauwaert et al., 2013). A2 virus-infected mice developed large VP1^+^ foci in the kidney cortex, but A2.V296F infected mice exhibited only small, sporadic VP1^+^ foci (**Figure 1C**). These results demonstrated that the V296F mutation impaired the ability of MuPyV to replicate in the kidney even under conditions of profound immunosuppression.

To determine whether the V296F mutation altered infection in the CNS, we infected mice intracranially (i.c.) and examined expression of LT mRNA. At 4 dpi following i.c. inoculation, A2.V296F showed equivalent infection in the brain to A2 (**Figure 1D**). To test if equivalent LT mRNA levels were indicative of infection in similar cell types, we examined brains 4 dpi for VP1^+^ cells by immunofluorescence microscopy. Infection with either virus resulted in sporadic VP1^+^ ependymal cells (vimentin^+^) or astrocytes (GFAP^+^) (**Figure 1E**) (Lavado and Oliver, 2011; Tissir et al., 2010). Because the S268F mutation is only seen in PML patients, we next asked whether A2 would outcompete A2.V296F *in vivo*. Mice received a 1:1 mixture (by PFU) of A2 and A2.V296F either i.c. or s.c.; the ratio of A2.V296F to A2 was determined at 14 dpi in various organs. To detect the relative levels of viral DNA, PCR primers were designed that amplified either a region of LT from both viruses or only the VP1 sequence of A2.V296F (**Figure S1C**). In mice infected s.c. a reduced A2.V296F:A2 ratio was seen in both the kidney and bone marrow, but this ratio was significantly higher and nearly equal in the brain (**Figure 1F**). In i.c.-inoculated mice A2.V296F infected the brain 1:1 with A2, but in the kidney was significantly outcompeted by A2, resulting in > 1:100 A2.V296F:A2 ratio in the kidneys of some mice (**Figure 1G**). These data indicated that compared to WT VP1, the V296F mutation caused decreased infection in the kidney, but equivalent infection in the brain.

We next asked whether CNS infection with A2.V296F caused similar encephalopathy as the A2 virus. Infection with either virus induced pronounced hydrocephalus of the lateral ventricles at 30 dpi (**Figure 2A**) with multiple foci of ablated ependyma and dysplastic changes in the choroid plexus (**Figure 2B**). Sham infected mice had single layer of vimentin^+^ cells adjacent to the ventricles, consistent with vimentin expression in the region being largely restricted to ependymal cells (**Figure 2C**) (Tissir et al., 2010). Brains of mice infected with either virus had expansion of the vimentin^+^ region abutting the ventricles, indicating damage to and/or disruption of the ependymal lining as a result of infection (**Figures 2C and 2D**). The vimentin^+^ cells in this region also had increased GFAP expression, possibly representing subventricular zone neural precursors responding to the disruption of the ependymal layer (**Figure 2C**) (Chojnacki et al., 2009). The ependyma and periventricular region had aggregates of CD3^+^ T cells and were diffusely infiltrated by Iba1^+^ cells (macrophages or microglia) in both A2- and A2.V296F-infected mice (**Figure 2E**). Together, these data demonstrated the V296F mutant virus retained the encephalogenic properties of the parental virus.

**Figure 2.**
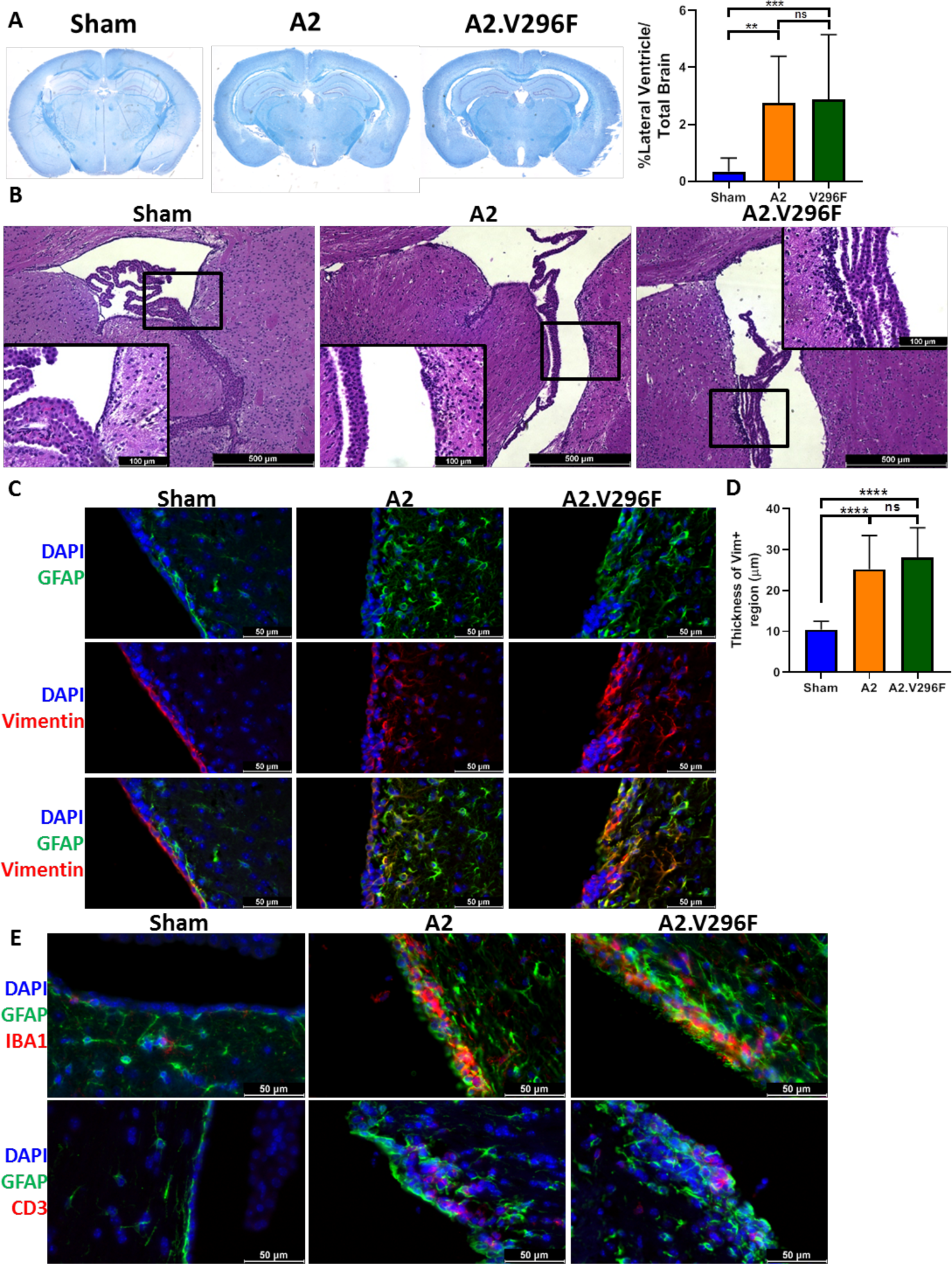
Persistent infection with either A2 or A2.V296F results in CNS pathology. (A) Left: LFB-PAS-stained brain sections 30 dpi with A2 or A2.V296F. Right: Hydrocephalus was quantified as the size of the lateral ventricle compared to total brain size. Data are from three independent experiments, n = 10-14 mice. (B) Representative 40x H&E images of the ependyma and choroid plexus of the lateral ventricle in mice 30 dpi. Inset image is 200x. (C) 400x fluorescence images of GFAP and vimentin expression in the lining of the lateral ventricles in mice 30 dpi. (D) Quantification of the thickness of the vimentin^+^ region shown in (C). Data are from three independent experiments, n = 13-15 mice. (E) 400x fluorescence images of Iba1^+^ and CD3^+^ cells in the lateral ventricles 30 dpi. Data were analyzed by one-way ANOVA (A, D). *p<0.05, ***p<0.001, ****p<0.0001.

### V296F confers resistance to a neutralizing VP1 antibody

Because this PML-like VP1 mutation in MuPyV was indistinguishable from the parental virus in CNS tropism and pathology, we explored the possibility that V296F allowed evasion of anti-polyomavirus humoral immunity, which is mediated by neutralizing VP1 antibody. Nearly all VP1 mutations in JCPyV-PML reside in one of the four solvent-exposed loops; these loops constitute the domains for binding the sialyated cell receptors and are the targets of the host’s antibody response (Buch et al., 2015; Lindner et al., 2019; Neu et al., 2010). Thus, we asked if A2.V296F was resistant to neutralization by the mAb 8A7H5 (Swimm et al., 2010). Incubation of MuPyV with 8A7H5 prior to infection potently neutralized A2 but did not affect infectivity by A2.V296F. Replacement of V296 with alanine did not abrogate neutralization by 8A7H5, indicating resistance to this mAb was mediated by certain amino acid mutations (phenylalanine) at the 296 position but not others (alanine) (**Figure 3A**). Spread of A2 infection in mouse fibroblast monolayers was significantly impaired by 8A7H5-containing media, but spread by A2.V296F was unimpeded (**Figure 3B**). To determine whether A2.V296F evaded neutralization by 8A7H5 *in vivo*, we passively immunized mice with 8A7H5 prior to s.c. infection and examined the efficacy of neutralization at day 4 or day 8 pi by measuring LT mRNA levels and the magnitude of the MuPyV-specific CD8 T cell response, respectively (**Figure 3C**). 8A7H5 immunization resulted in undetectable splenic LT mRNA levels in A2-infected mice 4 dpi, but had no effect on virus levels in A2.V296F-infected mice (**Figure 3D**); the identical pattern was seen for the anti-MuPyV CD8 T cell response (**Figure 3E**). Notably, we found that the IgG response to A2 MuPyV infection in WT mice was heavily biased toward the binding site recognized by 8A7H5, with 8A7H5 able to outcompete binding of over 80% of MuPyV-specific IgG to VP1 pentamers (**Figure 3F**). These findings suggested that a specific mutation in the sialic acid binding domains of VP1 diminished antibody neutralization of the virus.

**Figure 3.**
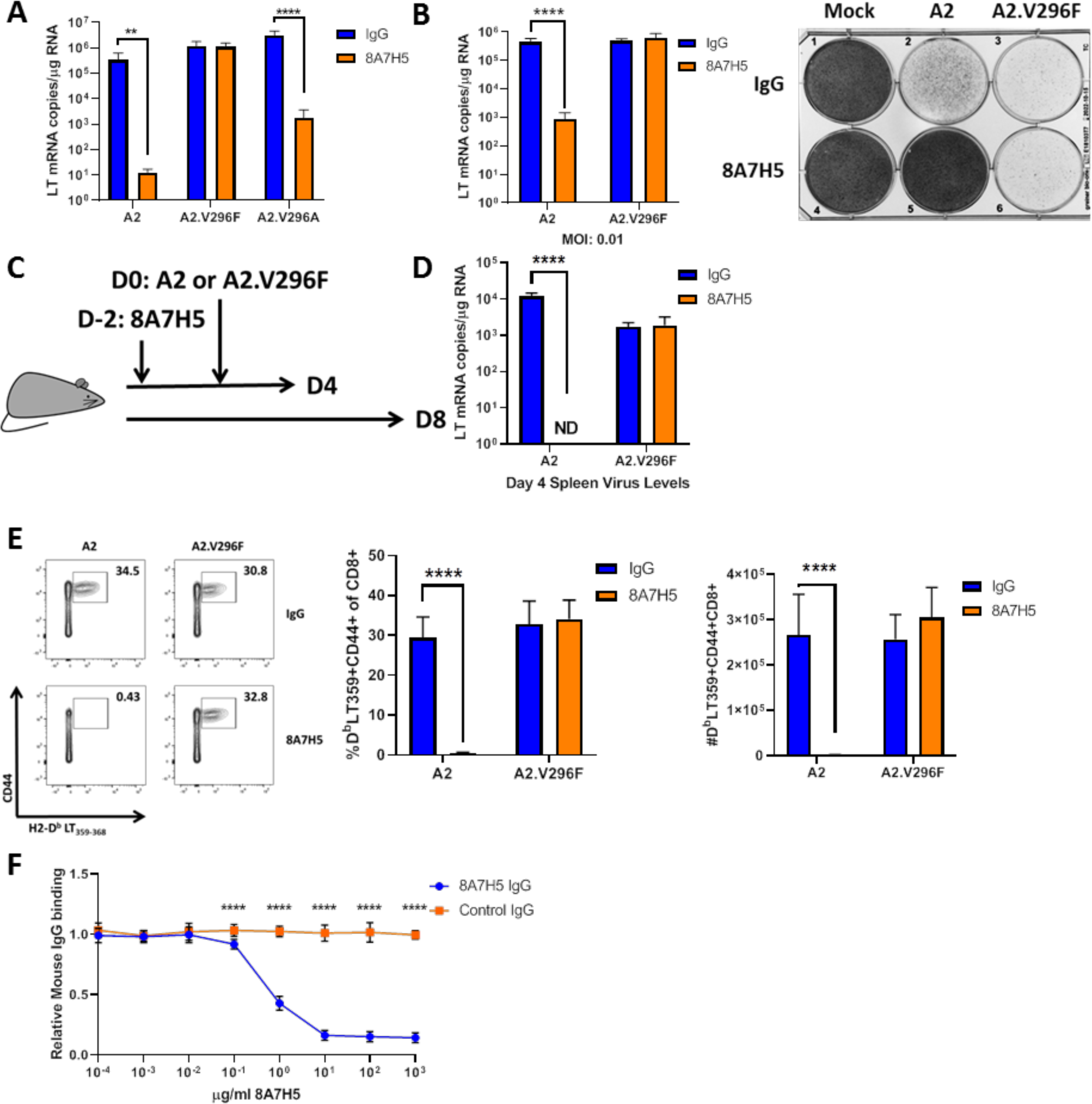
The V296F VP1 mutation confers resistance to a neutralizing mAb. (A) LT mRNA levels in NMuMG cells 24 h pi with A2, A2.V296A, or A2.V296F preincubated with 8A7H5 or control IgG. Data are from two independent experiments, n = 12. (B) Left: LT mRNA levels in A31 fibroblasts 96 hpi with A2 or A2.V296F at 0.01 MOI, 8A7H5 or control IgG was added to the media 24 hpi. Data are from two independent experiments, n = 6. Right: Protection from virus-induced cell death. A31 fibroblasts were treated as in left panel, fixed with formaldehyde and stained with crystal violet 7 dpi. (C) Experimental design for *in vivo* neutralization experiments. (D) LT mRNA levels in the spleens of mice injected with 8A7H5 or control IgG followed by infection with A2 or A2.V296F. Data are from two independent experiments, n = 6 mice. (E) Splenic T cell responses 8 dpi in mice treated and infected as in (D). Data are from two independent experiments, n = 6 mice. (F) Competition for binding to VP1 pentamers between immune sera and 8A7H5 IgG. Sera from mice 30 dpi with A2 was diluted to 2 μg/mL of VP1-specific IgG and combined with increasing concentrations of 8A7H5 or control IgG and measured for binding to VP1 pentamers by ELISA. Each sample was normalized to binding in the absence of exogenous IgG. Data are from three independent experiments, n = 12 mice. Data were analyzed by multiple t tests (A, B, D, E, F). **p<0.01, ***p<0.001, ****p<0.0001.

### Cryo EM reconstruction of MuPyV identifies mechanism of VP1 antibody escape by V296F

Similar to whole IgG, 8A7H5 Fab neutralizes and prevents spread of A2, but fails to do so for A2.V296F (**Figures S2A and S2B**). To investigate how V296F conferred resistance to 8A7H5, we collected cryo EM data for purified A2 capsids and A2 capsids incubated with saturating amounts of Fab (**Table S1**). Comparison of A2 and A2-Fab cryo EM micrographs revealed an obvious difference in particle size, demonstrating successful formation of virus:Fab complexes (**Figure 4A**).

**Figure 4.**
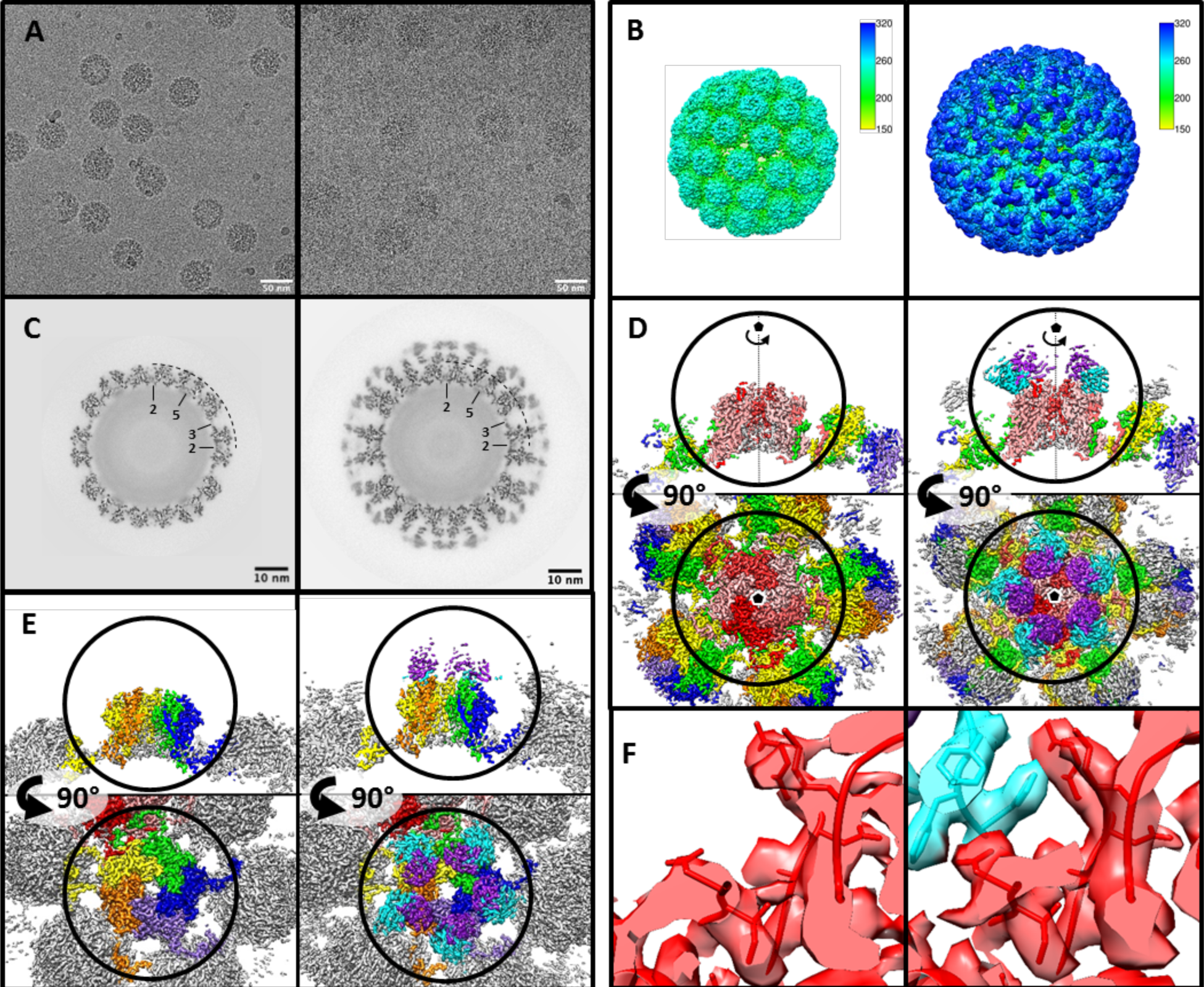
Cryo-EM image subvolume refinement reconstructions showing architecture of MuPyV-Fab complexes. (A) Micrographs of virus and virus-Fab complex (shown left and right, throughout figure) illustrate particle diameter difference due to bound Fab. (B) Surface rendered icosahedrally averaged maps. (C) Central sections demonstrate the quality of the maps and show the Fab and capsid densities are of comparable magnitude. (D & E) Subvolume refinement of pentavalent and hexavalent capsomers with sections through the maps (upper) and top-down views (lower) show the overall architecture. Pentavalent capsomers (D, VP1 density in shades of red) have fivefold symmetry (pentagon), whereas hexavalent capsomers have pseudo-symmetry (E, VP1 in OYGBV) most apparent in the contribution of VP1 C-terminal extensions to neighboring capsomers. Epitopes for the Fab molecules (light chain: purple; heavy chain: cyan) bridge adjacent VP1 molecules. (F) Local refinement of capsomer subvolumes resulted in interpretable sidechain density at the MuPyV-Fab interface (colors as in D&E).

Refinement of both datasets produced maps at 3.9 Å and 4.2 Å resolution for the A2 and A2-Fab complex, respectively (**Figure 4B and Table S2)** (Punjani et al., 2017; Zivanov et al., 2018). In contrast to the 60 epitopes present in the BKPyV:scFv structure (one per asymmetric unit), our complex map revealed density for 360 copies of 8A7H5Fab, corresponding to six copies per asymmetric unit, that is 1 Fab per each VP1 (**Figures S3A and S3B**) (Lindner et al., 2019). As a result, saturation with 8A7H5 Fab effectively blankets the entire surface of the virus (**Figure 4B**). The central section through the complex map revealed Fab density approximately equal to that of the capsid, indicating near saturation of the 360 available binding sites (**Figure 4C**).

After icosahedral refinement we next proceeded with local subparticle refinement of the constituent hexavalent and pentavalent capsomers for each dataset. This subparticle refinement allows each capsomer additional degrees of freedom to move independently of the rigid icosahedral matrix, which compensates for imperfect icosahedral symmetry present in flexible virus capsids (Goetschius et al., 2019; Stass et al., 2018). Using ISECC, our custom implementation of the localized reconstruction approach, we computationally generated subparticles from the refined whole particle images (**Figure S4**) (Ilca et al., 2015). Subvolume refinement improved the resolution of the pentavalent and hexavalent capsomers for both the A2 and A2-Fab complex (2.8 and 3.1 Å, respectively) (**Figures 4D and 4E**). For each dataset, the constituent capsomers attained approximately equal resolution (**Table S2**). This refinement also improved the resolution of the virus-Fab interface to 3.1-3.3 Å, with sidechain density clearly apparent (**Figures 4F, S5A, and S5B**).

Given the improved resolution attained by subvolume refinement, all atomic models were built in the subvolume maps (**Table S2**). Virus models were initialized with a VP1 (PDB ID 1SIE) and Fab structures (PDB ID 3GK8) mutated to match the primary structure of the A2 strain virus and 8A7H5 Fab. The Fab structure required manually rebuilding the complementarity determining regions (CDRs). All models were then refined into cryo EM density. Due to the quasi-equivalent VP1 molecules within the asymmetric unit (**Figure S3**), the six epitopes may have subtle conformational differences and during the build were not assumed to be identical. Therefore, one Fab was built into the pentavalent site and was subsequently docked into the remaining five hexavalent sites. After refining the build independently into each corresponding map density, loops comprising the six distinct epitopes in the asymmetric unit superimposed with a range of C alpha root mean square deviation (RMSD) of 0.40 Å to 0.51 Å. The resolution and map quality allowed us to identify unambiguously the 8A7H5 epitope and key contact residues (**Table 1**) for each of the six quasi-equivalent positions. Notably, these epitopes were found to be identical.

**Table 1.** VP1 contact residues within −0.4 Å van der Waal’s overlap. The conformational epitope spans three loops over two copies of VP1. Contributions from the adjacent VP1 are denoted with ‘.

| Loop | Residue |  |
| --- | --- | --- |
| BC | THR | 67 |
|  | GLU | 68 |
|  | ARG | 77 |
|  | GLY | 78 |
|  | ASN | 80 |
|  | THR | 83 |
|  | GLU | 91 |
| DE | PHE | 141` |
|  | LYS | 151` |
| HI | ARG | 292` |
|  | ASN | 293 |
|  | TYR | 294 |
|  | VAL | 296 |

The main contact residues of the Fab mapped to the heavy chain with minor contributions from the light chain (**Table S3**). The heavy chain made all contacts with one copy of the coat protein, whereas the light chain interacted with the adjacent VP1 (**Figure S5C**). On the virus capsid the 8A7H5 epitope consisted of thirteen residues of the BC and HI loops of VP1, and three residues from the DE (141, 151) and HI (292) loops of the adjacent VP1 (**Table 1, Figure 5A, and 5B**). Notably the 8A7H5 epitopes map directly adjacent to one another in a ring tracing around the contours of the capsomer. There was a predicted salt bridge between VP1 R77 and D99 located in CDR loop H3 of 8A7H5 (**Figure 5C**).

**Figure 5.**
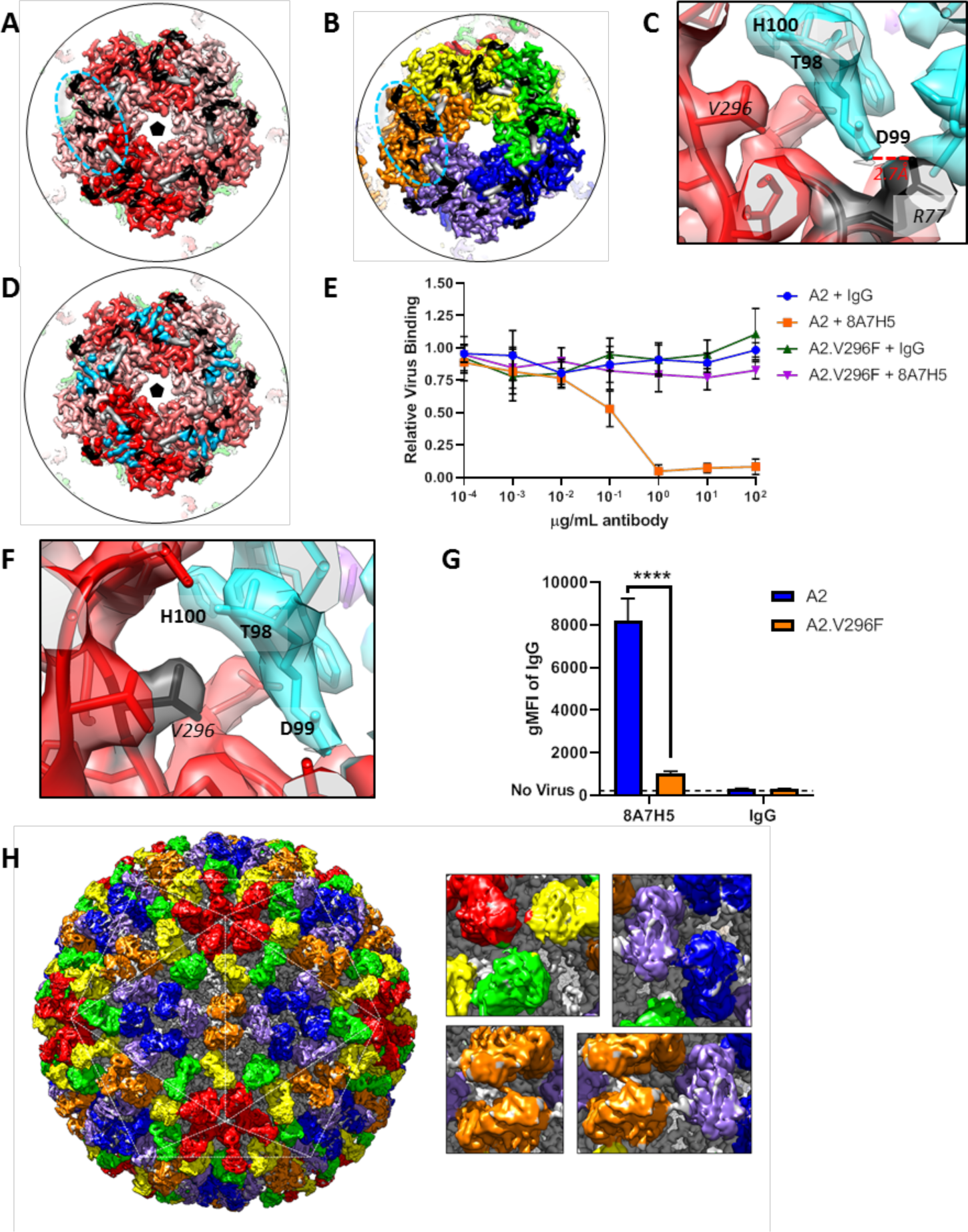
Cryo EM reconstruction of MuPyV identifies mechanism of VP1 antibody escape by the V296F mutaton. (A & B) The Fab epitope bridges adjacent copies of VP1 on the pentavalent capsomer (A, shades of red) and hexavalent capsomer (B, OYGBV). Neighboring epitopes abut directly against each other. Contact residues from the main VP1 chain are noted in black, with minor contributions from the adjacent VP1 in grey. (C) A salt-bridge is formed between R77 and Fab heavy chain residue D99. This interaction is near key residue V296, despite the large distance in linear sequence. (D) The Fab epitope and receptor binding residues (sky blue) overlap (PDB ID 5CPY) (Buch et al., 2015). (E) Increasing concentrations of 8A7H5 prevent the attachment of A2, but not A2.V296F. H2B-GFP labeled virus was incubated with antibody prior to incubation with NMuMG cells. GFP fluorescence was measured by flow cytometry. Data are from two independent experiments, n = 6. (F) The V296F mutation would place a bulky residue at the MuPyV-Fab interface, disrupting the Fab heavy chain residue T98, D99, and H100 (cyan) interactions. (G) V296F prevents the binding of 8A7H5 to VP1. NMuMG cells were incubated with A2 or A2.V296F followed by incubation with 8A7H5. Bound 8A7H5 was detected with an anti-IgG secondary. Data are from two independent experiments, n = 6. (H) Six quasi-equivalent Fab molecules (ROYGBV) are contained within the asymmetric unit without clashes, despite the close proximity of Fab constant domains (inset). Data were analyzed by Mann-Whitney U test (E). ****p<0.0001.

The 8A7H5 epitope overlapped with residues associated with receptor binding (**Figure 5D**). This overlap suggests the mechanism of antibody neutralization is to block the receptor binding site and prevent virus interaction with the host cell. To address this experimentally, we analyzed the binding of H2B-GFP labeled A2 or A2.V296F to cells after pre-incubation of virus with 8A7H5. 8A7H5 mAb/Fab blocked A2 viral attachment, but had no effect on A2.V296F attachment assayed by flow cytometry (**Figures 5E and S2C**). The V296F mutation placed a bulky phenylalanine sidechain directly within the Fab-virus interface likely disrupting the interaction with the CDR loop H3 through steric hinderance (**Figure 5F**). Consistent with this model, direct binding assays showed a significant reduction in 8A7H5 binding to A2.V296F compared to A2 (**Figures 5G and S2D**). The adjacent placement of epitopes resulted in a striking and tightly packed arrangement of bound Fab both within a capsomer (variable domain) and between capsomers (constant domain). Surprisingly there was no steric clash observed between Fabs bound to neighboring epitopes (**Figures 5H and S5D**). This observation indicates that the MuPyV capsid was able to accommodate 360 copies of 8A7H5 Fab, saturating all available epitopes.

### Additional PML mutations in MuPyV impair kidney infection and disrupt 8A7H5 binding

Comparing the VP1 structures of JCPyV and MuPyV, we identified several additional substitutions to introduce into MuPyV to mimic other PML mutations (Gorelik et al., 2011; Sunyaev et al., 2009) (**Figure 6A**). Of these mutations, only those at N293 and V296 resulted in viable MuPyV variants. All mutant viruses showed a defect in kidney infection following s.c. inoculation, similar to A2.V296F (**Figure 6B**). Inoculation i.c. with these viruses resulted in similar levels of infection in the brain compared to A2 (**Figure 6C**). The mutant viruses, however, varied in their susceptibility to 8A7H5-mediated neutralization. Predictably, V296Y conferred complete resistance to 8A7H5. A2.N293F and A2.N293Y remained sensitive to 8A7H5 neutralization (**Figure 6D**).

**Figure 6.**
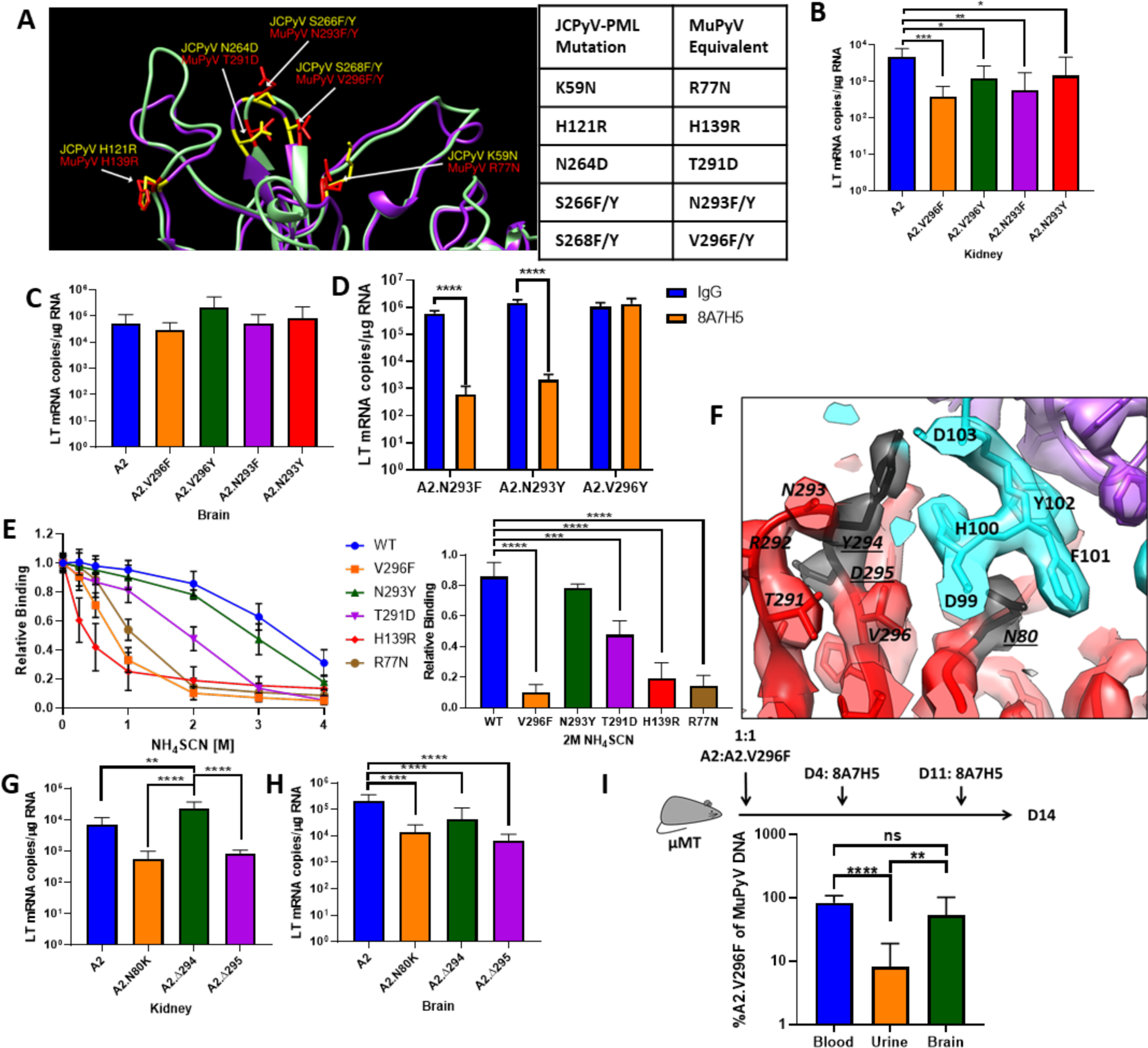
Additional JCPyV-PML mutations in MuPyV impair kidney, but not brain infection, and have varying effects on VP1 mAb neutralization. (A) Structural comparison of PML mutation sites in JCPyV VP1 (PDB 3NXG) with MuPyV VP1 (PDB 5CPU) residues (Buch et al., 2015; Neu et al., 2010). (B) Kidney LT mRNA levels in mice 4 dpi with A2 or mutant viruses s.c. Data are from two independent experiments, n = 9-10 mice. (C) Brain LT mRNA levels in mice 4 dpi with A2 or mutant viruses i.c. Data are from three independent experiments, n = 9-10 mice. (D) LT mRNA levels in NMuMG cells 24 hpi with A2.V296Y, A2.N293F, A2.N293Y preincubated with 8A7H5 or control IgG. Data are from two independent experiments, n = 12. (E) Analysis of 8A7H5 avidity for mutant VP1’s using NH_4_SCN. Left: Relative binding of 8A7H5 to recombinant VP1 with increasing concentrations of NH_4_SCN. Right: Relative binding of 8A7H5 to recombinant VP1 at 2M NH_4_SCN. Each point is the average of two technical replicates in an independent repeat, n = 3. (F) Location of novel changes in the VP1 escape mutations. (G) LT mRNA levels 4 dpi in the kidneys of mice infected s.c. Data are from two independent experiments, n = 8 mice. (H) LT mRNA levels 4 dpi in the brains of mice infected i.c. Data is from 2-4 independent experiments, n = 8-16 mice. (I) Viral shedding in the urine is impaired by V296F, despite antibody escape in the blood. Top: Experimental design for infection of μMT mice and 8A7H5 treatment. Bottom: Frequency of A2.V296F DNA in blood, urine, and brain tissue of mice. Data are from three independent experiments, n = 13-14 mice. Data were analyzed by one-way ANOVA (B, C, E Right, G, H, I) or multiple t tests (D). In E, comparisons

To examine the effect of the VP1 mutations that failed to produce virus, we generated recombinant VP1 pentamers with these mutations to test 8A7H5 binding. We further measured 8A7H5 avidity for VP1 by ELISA combined with NH_4_SCN treatment, a chaotropic agent that disrupts low-affinity antibody interactions (Pullen et al., 1986) (**Figure 6E**). 8A7H5 showed high avidity interactions with WT and N293Y VP1 pentamers, and low avidity with V296F pentamers. The R77N and H139R mutations each reduced 8A7H5 avidity to a similar extent as V296F, suggesting that these mutations would also confer resistance to 8A7H5. The T291D mutation, also decreased 8A7H5 avidity, but not to the extent of R77N, H139R, and V296F (**Figures S6A and S6B**). Collectively, these data demonstrated that impaired kidney and retained brain tropism is a common theme for several PML mutations, but only a subset of these different mutations are capable of evading recognition by this VP1 mAb.

### 8A7H5 mAb selects VP1 escape mutations

We serially passaged A2 in the presence of 8A7H5 to select for *de novo* escape mutants. Three VP1 mutant viruses were isolated, an N80K point mutation in the VP1 BC loop and two single amino acid deletions, Δ294 and Δ295, adjacent to V296F in the HI loop. These mutations each conferred complete resistance to 8A7H5-mediated neutralization (**Figure S6C**). VP1 residues N80, Y294, and D295 each map to or near contact residues predicted by the virus-Fab complex structure (**Figure 6F and Table 1).** Infection of mice s.c. with the mutant viruses resulted in elevated virus levels of A2.Δ294 and reduced virus levels of A2.N80K and A2.Δ295 in the kidney, indicating these mutations have varying effects on kidney tropism (**Figure 6G**). All three mutant viruses had reduced brain infection levels following i.c. inoculation compared to A2 (**Figure 6H**). These differences in kidney tropism and decrease in brain tropism indicated that although these mutations shared 8A7H5-resistance with A2.V296F, individual mutations in this region had varying effects on tropism in an organ-specific manner.

### A2.V296F shows poor shedding under conditions of antibody escape

Our data suggested a model where PML-associated VP1 mutations promote antibody escape at the expense of infection and persistence in the kidney. This predicts that A2.V296F would be poorly shed in the urine, even under antibody escape conditions in the host. μMT mice have a genetic defect in B cell development and fail to mount an anti-MuPyV antibody response (Kitamura et al., 1991; Szomolanyi-Tsuda and Welsh, 1996). To approximate the clinical observation of WT virus in the urine of PML patients despite mutant virus being in the blood and CSF, we inoculated μMT mice with a 1:1 ratio of A2:A2.V296F and began administering 8A7H5 4 dpi. At 14 dpi, virus shed in the urine was heavily biased towards A2, despite the mice having high levels of A2.V296F in the blood and brain tissue (**Figure 6I**). This result showed that a severe impediment to kidney replication limits shedding of the V296F mutant virus in the urine, despite being viremic.

## DISCUSSION

In this study, we elucidated the impact of MuPyV VP1 mutations on viral tropism and antibody neutralization, drawing a mechanistic link between JCPyV capsid mutations and PML pathogenesis. We applied a custom subvolume refinement approach to reconstruct cryo EM images of native capsid:antibody complexes at atomic resolution. The structures revealed the mechanism of VP1 antibody evasion. Using MuPyV with a VP1 mutation matching a frequent VP1 mutation in JCPyV-PML, we found that this viral variant retained tropism for the CNS, but was profoundly impaired in its ability to replicate in the kidney, a major organ reservoir for persistent polyomavirus infections. This mutation blocked neutralization by a MuPyV VP1 mAb via steric hindrance. Other JCPyV-PML VP1 mutations introduced into MuPyV also impaired kidney, but not brain infection, and varied in their ability to bind the VP1 mAb. mAb-escape MuPyV variants selected *in vitro* used additional mechanisms to evade neutralization but exhibited altered replication in both the brain and kidney. This disconnect between nAb escape and CNS tropism shows that only a subset of JCPyV VP1 variants refractory to the VP1 antibody response are detected in PML patients. By extension, our data support the concept that the VP1 antibody response selects JCPyV variants capable of causing CNS injury.

Our implementation of subvolume refinement allowed the rapid solution of the highest resolution cryo EM structures of any polyomavirus map to date. This innovation was achieved using fewer particles compared to the traditional cryo EM approach **(Table S4**). Improvements seen after subvolume refinement may be attributable to capsid flexibility and the defocus gradient extending over the 45 nm capsid. Curiously, correction for optics aberrations and the Ewald sphere effect improved resolution of only the icosahedrally averaged MuPyV capsid map, but not the MuPyV-Fab complex map (Zivanov et al., 2020). Resolution improvement from subvolume refinement of MuPyV was equivalent to that seen with optics refinement, but these improvements were not cumulative. In contrast, the MuPyV-Fab complex map improved after subvolume refinement. The lack of improvement from Zernicke parameter refinement for the complex map may be because refinement of optics parameters is a reference-based process, such that the flexibility contributed by 360 copies of Fab may be a barrier to properly solving and correcting for optical aberrations.

The densely packed arrangement of 360 Fab molecules coating the capsid (**Figure 5H**) signifies the presence of six structurally identical 8A7H5 epitopes within the asymmetric unit, despite the quasi-equivalence of VP1 molecules that form pentavalent and hexavalent capsomers. The conformational epitope bridges adjacent VP1 molecules within each capsomer, yet without provoking clash between neighboring Fab molecules. This binding pattern sharply contrasts with scFv 41F17, which recognizes a single structurally unique epitope within the BKPyV asymmetric unit (Lindner et al., 2019). The difference in binding behavior is due to the apical location of the 8A7H5 epitope that is comprised of structural features common to all capsomers. In contrast, the 41F17 epitope is laterally located and formed through the interaction of VP1 chains between adjacent hexavalent capsomers.

The structural data also explain antibody escape caused by VP1 mutations in JCPyV-PML when mapped into MuPyV. Introduction of bulky F/Y sidechains at position 296 within the antibody footprint likely promotes escape through steric collision, since an A residue at 296 retains neutralization. Loss of antibody binding to R77N pentamers is probably because of the lost salt bridge between VP1 R77 and D99 in CDR loop H3 of 8A7H5. T291D had an intermediate effect on 8A7H5 binding, consistent with its location immediately adjacent to several crucial contacts in the HI loop. Because H139 is not directly in the footprint, the mechanism of lost binding is not readily apparent but may be due to a long distance interaction. Although N293 (corresponding to JCPyV S266) is identified as a contact residue, it is on the periphery of the 8A7H5 footprint, which may explain retained recognition and neutralization of N293F/Y by 8A7H5. A recent report showed that sera from healthy individuals and JCPyV sero-positive patients failed to recognize S266F but not wild type (Jelcic et al., 2015). Additionally, broadly neutralizing scFv 41F17 recognized an epitope comprised of residues from VP1 proteins between hexavalent capsomers (Lindner et al., 2019). Thus, mutations may disrupt recognition by antibodies with epitopes distinct from that of 8A7H5.

Our cryo EM complex structures also explain three spontaneous escape mutations found during serial passage in the presence of 8A7H5. Deletion of contact residue Y294 or the immediately adjacent D295 provide escape through shortening and reorganization of the key antigenic HI loop. N80 directly interacts with 8A7H5 via a trio of residues (H100, F101, Y102) in CDR loop H3 of 8A7H5 that form a depression in the Fab topology into which the N80 side chain inserts (**Figure 6F**). The N80K substitution would disrupt this interaction by introducing a positive charge and a longer sidechain.

Several factors may lead to the strong neutralizing activity of 8A7H5 mAb. Because the 8A7H5 epitope bridges neighboring VP1 molecules within each capsomer, 8A7H5 binding may stabilize the virus and prevent uncoating. A salt-bridge is formed between the virus and antibody, strengthening the interaction. The conformational epitope and the receptor binding site both contain the HI loop; this significant overlap allows antibody binding to prevent attachment to the cellular receptor. 8A7H5 Fab recognizes 360 structurally identical epitopes on the virus capsid, despite the quasi-equivalence of the six VP1 chains within the asymmetric unit. There is no steric clash between the 360 copies of 8A7H5 Fab, resulting in the striking and tightly packed arrangement of Fab seen in **Figure 5H.** It is important to note that this packing would be unlikely to occur *in vivo* due to the bulk of a whole antibody.

JCPyV-PML VP1 mutations have been proposed to drive neurovirulence or evasion of humoral immunity, but not both (Geoghegan et al., 2017; Jelcic et al., 2015; Maginnis et al., 2013; O’Hara et al., 2018; Ray et al., 2015). Our data reconcile these findings, indicating these mutations impair infection in sites of typical polyomavirus persistence (kidney, bone marrow), but retain infectivity in the CNS. We demonstrated that a MuPyV with the V296F PML-like VP1 mutation had profoundly impaired kidney tropism and lower viruria than parental MuPyV. Likewise, only archetype JCPyV is detected in urine, whereas VP1 mutant viruses are found in blood and CSF (Gorelik et al., 2011; Reid et al., 2011). This impaired kidney tropism by PML-VP1 mutants may underlie the absence of JCPyV-associated nephritis in PML patients, despite the kidney being the major site of JCPyV persistence (Berger et al., 2017).

Emergence of VP1 mutant JCPyVs in PML patients but not healthy individuals infers that viruses with these mutations have a replication advantage in the setting of depressed immune status (Zheng et al., 2005). Mutations in the four receptor-binding loops of VP1 are typically detrimental to viral fitness and persistence (Bauer et al., 1999; Caruso et al., 2003), and our data showing altered tropism by MuPyV VP1 mutants are clearly aligned with this idea. Evasion of host antibodies provides a strong selective pressure to promote the spread of an otherwise replication-disadvantageous mutation. Experimental demonstration of this scenario comes from evidence that the mutant A2.V296F virus strongly outcompeted parental A2 virus in the blood and brain, but was still poorly shed into the urine, when faced with an A2-nAb (**Figure 6I**). Our data agree with recent reports showing poor neutralization by sera from PML patients for their VP1 mutant JCPyVs, and indicate that selection of VP1 mutants is driven by an antibody response sufficient to control parental but not a VP1 mutant virus (Jelcic et al., 2015; Ray et al., 2015). Thus, our findings indicate that JCPyV takes a hit to viral fitness in order to evade humoral immunity.

By extension, our results strongly support the concept that antibody escape is a requisite first step in PML development. The resulting viremia, then, would precede viral entry into the brain, whether by infiltrating the CSF via the choroid plexus, direct infection of brain endothelium, or by hitchhiking a cellular vehicle (Chapagain et al., 2007; Dörries et al., 2003; Houff et al., 1988; von Einsiedel et al., 2004). JCPyV viremia is found in multiple sclerosis patients treated with natalizumab (Major et al., 2013). In support of a choroid plexus-mediated route, JCPyV infects primary choroid plexus epithelial cells, and JCPyV-infected choroid plexi are found in PML brains (Corbridge et al., 2019; O’Hara et al., 2020, 2018). Both A2 and A2.V296F viruses productively infect the ependyma, and we reported ependymal infection by MuPyV under conditions of immune suppression (Mockus et al., 2020). Infection of the choroid plexus and ependyma may serve as a viral staging area for JCPyV invasion of the brain parenchyma, providing a foothold for viral dissemination in the CNS parenchyma.

Using the MuPyV CNS infection model, we demonstrate that evasion of the host’s neutralizing antiviral humoral response is the dominant driver of VP1 mutant viruses that retain CNS tropism. We developed a custom subvolume refinement approach to reconstruct efficiently cryo EM structures of polyomavirus capsid-Fab complexes at the highest resolution to date. These structures elucidated the mechanisms of neutralization and antibody escape. Our findings argue against the concept that VP1 mutations act per se to render JCPyV neurovirulent. Instead our work supports the model that viremia, consequent to outgrowth of antibody-escape VP1 variants, is a critical step in PML pathogenesis.

## Supporting information

Key Resources Table

## Acknowledgments

The authors thank the staff of the Penn State College of Medicine Flow Cytometry Core Facilty, N. Sheaffer, J. Bednarczyk, J. Vogel, and J. Zhang for assistance with flow cytometry analysis; G. Snavely and E. Mullady of the Comparative Medicine Histology Core for the preparation of tissue sections; B. Garcea for the generous gift of the PyVP1-pGEX-4T-2 plasmid and rabbit VP1 antisera; and the staff of the Department of Comparative Medicine at the Penn State College of Medicine. Research reported in this publication was supported by the National Institute of Neurological Disorders and Stroke, the National Institute of Allergy and Infectious Diseases, and the National Cancer Institute under award numbers R01NS088367 and R01NS092662 (AEL), R01AI107121 (SLH), F32NS106730 (CSNW), F31NS083336 (ELF), and T32 CA60395 (MDL and SMB), and the Jake Gittlen Laboratories for Cancer Research (NDC). The content is solely the responsibility of the authors and does not necessarily represent the official views of the National Institutes of Health. In addition, support for this work was provided by Pennsylvania Department of Health CURE funds (SLH).

## Author Contributions

MDL, DJG, SLH, and AEL conceived the study. MDL, CSNW, KNA, GJ, DGH, and ELF conducted the cell culture, biochemistry, ELISAs, immunohistochemistry, virology, and mouse experiments. SMB and NDC sequenced the mAb. SHC prepared sample for cryo EM data collection. CMB collected the cryo EM data. DJG designed and developed the custom software, solved the structures, and built the models. MDL, DJG, CSNW, SLH, and AEL interpreted the data. MDL, DJG, SLH, and AEL wrote the manuscript.

## MATERIALS and METHODS

Additional details on reagents and resources are listed in the Key Resources Table.

### Mice

C57BL/6 mice were purchased from the National Cancer Institute and μMt mice were purchase from the Jackson Laboratories. STAT1^-/-^ mice (The Jackson Laboratory) were kindly provided by Dr. Christopher Norbury (Penn State College of Medicine). Male and female mice were used for experiments between 6-15 weeks of age. Mice of the same sex/age were randomly assigned to experimental groups. Mice were housed and bred in accordance with the National Institutes of Health and AAALAC International Regulations. The Penn State College of Medicine Institutional Animal Care and Use Committee approved all experiments.

### Virus strains

All work was performed with the A2 strain of MuPyV. Viral stocks were generated by transfection of viral DNA into NMuMG cells using Lipofectamine™ 2000 Transfection Reagent (ThermoFisher). A single passage in NMuMG cells was used for viral amplification to generate a high titer virus stock. Virus stocks were titered on A31 fibroblasts by plaque assay (Lukacher and Wilson, 1998).

### Cell lines and primary cells

The 8A7H5 hybridoma was previously generated by immunization of a rat with MuPyV VP1 virus-like particles (Swimm et al., 2010). NMuMG, BALB/3T3 clone A31 “A31”, and mIMCD-3 cells were purchased from ATCC. Mouse embryonic fibroblasts (MEFs) were isolated from day 13 C57BL/6 embryos. The 8A7H5 hybridoma was maintained in PFHM-II Protein-Free Hybridoma Medium (ThermoFisher) at 37°C in 5% CO_2_. mAb was generated by growing the hybridoma in a CELLine disposable bioreactor flask (Corning). All other cells were maintained in Dulbecco’s Minimal Eagle Media supplemented with 10% fetal bovine serum, 100 U/mL penicillin, and 100 U/mL streptomycin (DMEM) at 37°C in 5% CO_2_. The sex of NMuMG cells is female, the sex of mIMCD-3 and A31 cells is not reported. Cell lines were not specifically authenticated, but were examined for correct cell morphology and used at low passage.

### Generation of mutant viruses

Viral mutants were generated by site-directed mutagenesis using the Quikchange II Site-directed mutagenesis kit (Agilent) with forward and reverse primers specific for each mutation (Key Resources Table). To confirm the presence of the mutations, viral DNA was isolated from virus stocks and the VP1 region was PCR amplified and sequenced.

### Virus infections

Mice were infected with MuPyV s.c. via the hind footpad with 1×10^6^ PFU or i.c. with 5×10^5^ PFU. For *in vivo* neutralization experiments, mice received 250μg of 8A7H5 or control IgG in PBS intraperitoneally on the specified days. For *in vitro* experiments, subconfluent cells were incubated with virus for 1.5 h at 4°C, and then free virus was removed by washing with DMEM. For single cycle replication and plaque assays, free virus was not removed. Following infection, cells were maintained in DMEM at 37°C in 5% CO_2_.

### Viral genome quantification

50 μL of viral lysate was treated with 250 U of Benzonase® Nuclease (Sigma) in 250 μM MgCl_2_ at 37°C for 1 h. Viral genomes were then isolated using the Invitrogen Purelink Viral RNA/DNA Mini Kit (ThermoFisher Scientific). Viral genomes were quantified by Taqman qPCR with primers and probe targeted to the LT region of the viral genome (Key Resources Table) (Wilson et al., 2012).

### Infection neutralization assay

10 μg of 8A7H5 mAb/Fab or control IgG/Fab (Jackson ImmunoResearch) was incubated at 4°C for 30 minutes with 1×10^4^ PFU of MuPyV and then added to 1×10^5^ NMuMG cells. Cells were infected at 4°C for 1.5 h and mRNA was harvested 24 h later.

### Viral mRNA quantification

RNA was harvested with TRIzol Reagent (ThermoFisher) and isolated by phenol:chloroform extraction followed by isopropanol precipitation. cDNA was prepared with 1-2 ug of RNA using random hexamers and Revertaid RT (ThermoFisher). LT mRNA levels were quantified by Taqman qPCR with normalization to TATA-Box Binding Protein and compared to a standard curve (Maru et al., 2017).

### H2B-GFP labeling of MuPyV and 8A7H5 mAb attachment assays

Virus was labeled by infection of NMuMG cells expressing an H2B-GFP fusion protein, which is incorporated into the PyV minichromosome during DNA replication and packaging (Fang et al., 2010; Geoghegan et al., 2017; Kanda et al., 1998). 8A7H5 mAb or control IgG was incubated with labeled A2 or A2.V296F at a ratio of 5,000 encapsidated viral genomes/cell for 30 m at 4°C, then added to a suspension of 5×10^4^ trypsinized NMuMG cells and incubated for 30 m at 4°C. Cells were then washed twice in PBS and fixed for 20 m in 2% PFA. GFP fluorescence on the cells was quantified using a BD LSRFortessa Flow Cytometer and normalized to fluorescence of virus bound in the absence of antibody. To measure 8A7H5 attachment to virions, virus was incubated with cells for 30 m at 4°C, then treated with 8A7H5. Bound 8A7H5 was stained with APC anti-rat IgG and quantified using a BD LSRFortessa Flow Cytometer.

### Fab generation and mAb sequencing

8A7H5 and control Fabs were generated using the Pierce Fab Micro Preparation Kit (Thermo Fisher) and purified on Protein G columns (Thermo Fisher). Sequencing of the heavy and light chains of the mAb was carried out as previously described (Guan et al., 2015). In brief, hybridoma cells were pelleted and RNA was extracted with TRIzol Reagant (ThermoFisher). cDNA was generated with Revertaid RT (ThermoFisher) and amplified by PCR using *PfuTurbo* DNA Polymerase (Agilent) with published primers (Wang et al., 2000). PCR products were purified using the QIAquick PCR Purification Kit (Qiagen) and sequenced.

### Virus purification for Cryo EM

Virus purification was adapted from a published BKPyV purification method (Hurdiss et al., 2018). NMuMG cells were infected at low MOI with A2 or A2.V296F. Following cell lysis, media/lysate was collected and cell debris was pelleted at 15,000g for 20 m. The supernatant was collected, and the pellet was resuspended in Buffer A (10mM Hepes, 1mM CaCl_2_, 1mM MgCl_2_, 5 mM KCl). The pellet was freeze-thawed 3 times, treated with Benzonase® Nuclease (75 U/mL) (Sigma) and type II neuraminidase (1/2000) (Sigma) for 1 h at 37°C, then combined with 0.1% deoxycholic acid and incubated at 37°C for 15 m, followed by 42°C for 5 m. The sample was pelleted at 15,000g for 20 m, and the supernatant was collected and combined with the original supernatant. The combined supernatants were layered on a 20% sucrose cushion and spun for 3 h at 85,000g. The pellet was resuspended in Buffer A and layered on top of a 27/33/39% gradient of OptiPrep (STEMCELL) in Buffer A. The sample was spun at 237,000g for 3.5 h at 16°C. The band containing the virus was then removed with a syringe.

### VP1 pentamers

Full-length MuPyV VP1 in the pGEX-4T-2 expression plasmid was provided by Robert Garcea (University of Colorado, Boulder). VP1 mutants were generated using the Quikchange II Site-directed mutagenesis kit (Agilent) with forward and reverse primers listed in the Key Resources Table. VP1 pentamers were induced by IPTG in BL21 *E. coli* (Agilent), and purified with glutathione sepharose (GE Healthcare) followed by thrombin cleavage. Pentamers were then bound and eluted from a cellulose phosphate column.

### ELISA

ELISAs were performed using 50 ng of VP1 pentamer/well in an EIA/RIA Polystyrene High Bind Microplate (Fisher Scientific) coated overnight at 4°C. For 8A7H5 competition with immune mouse sera, the VP1-specific IgG concentration of the serum was measured, then 100 ng of VP1-specific IgG was combined with increasing concentrations of 8A7H5 or control IgG for the ELISA. Bound mouse IgG was detected with a mouse IgG-specific secondary (Biolegend). For avidity measurements, 8A7H5-pentamer complexes were treated with NH_4_SCN in 0.1 M phosphate for 15 m before detection of 8A7H5 mAb. For each VP1 variant, 8A7H5 binding was normalized to signal in the absence of NH_4_SCN.

### Flow cytometry

Single cell suspensions of splenocytes were stained with antibody/tetramer cocktails (see Key Resources Table) in 100 μL for 30 m and quantified using a BD LSRFortessa Flow Cytometer. Flow cytometry data were analyzed using FlowJo software (BD).

### mAb-mediated selection for VP1 escape mutants

1×10^5^ NMuMG cells were infected with A2 at an MOI of 0.1. 24 hpi, 0.5 μg/mL 8A7H5 was added to the media. The media was replaced every 3-4 days until cell death 1-2 weeks post infection. The lysate was collected and diluted 1/100 to infect new NMuMG cells. After 3-4 passages, the resulting lysate was diluted 1/100 and combined with 10 μg of 8A7H5 for 30 m prior to infection of NMuMG cells. Following infection, the cells were maintained in 10 μg/mL 8A7H5 and observed for cell death/lysis. Lysates were then collected and viral DNA was isolated using the PureLink® Viral RNA/DNA mini Kit (ThermoFisher). The VP1 region of the genome was then amplified by PCR and sequenced. Identified mutants were cloned or generated by site-directed mutagenesis and confirmed to be escape mutants by neutralization assay.

### Immunofluorescence microscopy and histology

Mice were perfused with 10% heparin in PBS, followed by 10% neutral buffered formalin (NBF). Heads were fixed overnight in NBF and brains were then removed, paraffin-embedded, and sectioned. For histology, sections deparaffinized and were stained with H&E or Luxol Fast Blue-Periodic Acid Schiff (LFB-PAS). For immunofluorescence, sections were deparaffinized and subjected to antigen retrieval (95°C 10 mM sodium citrate pH 6 for 10 m). Sections were permeabilized with 1% TritonX-100 for 15 m, washed 2x in PBST (0.1% Triton X-100, 0.05% Tween20), then blocked with blocking buffer (5% BSA in PBST) and incubated overnight at 4°C with primary antibodies in blocking buffer. Sections were washed 3x with PBST, incubated with secondary antibodies in blocking buffer, washed 3x with PBST, then mounted with ProLong Gold Antifade with DAPI (Thermofisher). Images were acquired on a Keyence BZ-X710 all-in-one fluorescence microscope or a Leica DM4000 fluorescent microscope for histology or immunofluorescence, respectively. The thickness of the vimentin^+^ region was quantified across six images/sample in ImageJ (NIH). Hydrocephalus was quantified by measuring the pixel area of right and left lateral ventricles and dividing by the pixel area of the total brain section (Mockus et al., 2020). For representative fluorescence and brightfield images, adjustments for brightness/contrast were done uniformly to all images in the group using LAS X (Leica).

### DNA isolation and quantification

Solid tissues were homogenized using a TissueLyser II (Qiagen). DNA was isolated using the Wizard® Genomic DNA Purification Kit (Promega). DNA was isolated from lysates, blood, and urine using the PureLink® Viral RNA/DNA mini Kit (ThermoFisher). For competition experiments, total viral DNA and V296F DNA was quantified by Sybr Green qPCR (Quantabio) using the LT DNA and V296F DNA qPCR primers, respectively (Key Resources Table). The ratio of A2:A2.V296F was determined by comparison to a standard curve of known A2:A2.V296F DNA ratios.

### Cryo EM and Data Collection

MuPyV was buffer exchanged against 10 mM HEPES pH 7.9, 1 mM CaCl_2_, 1 mM MgCl_2_, 5 mM KCl (Hurdiss et al., 2016). MuPyV (2.8 mg/mL) was incubated with 8A7H5 Fab (1.1 mg/mL) for 30 m at room temperature. For vitrification of each sample, a 3 µL aliquot was applied to a freshly glow-discharged QUANTIFOIL EM grid. Grids were blotted for 3 s in 95% relative humidity before plunging into (Vitrobot; Thermo Fisher) liquid ethane. Cryo-EM datasets were collected at 300 kV with a Titan Krios microscope (Thermo Fisher) equipped with a spherical aberration corrector at the Huck Institute for Life Sciences cryo EM Facility. Automated single-particle data acquisition was performed with EPU using the Falcon 3 detector with a nominal magnification of 59,000x, yielding a final pixel size of 1.1 Å/pixel (**Table S2**).

### Image processing

Patch motion correction, patch-CTF estimation, particle picking, 2D classification, and icosahedral refinement were performed in cryoSPARC (Punjani et al., 2017). Particles transferred to RELION version 3 for polishing, before another round of icosahedral refinement (Asarnow et al., 2019; Zivanov et al., 2018). Pentavalent and hexavalent subparticles were extracted using ISECC, our custom implementation of the localized reconstruction technique [(Ilca et al., 2015). ISECC was written for compatibility with RELION 3.1, with additional features for correlative subparticle analysis (code available at https://git.psu.edu/suh21/isecc). Subparticles were locally refined in RELION.

### Model building

All models were built into the corresponding subparticle maps, rather than icosahedral maps. VP1 models were initialized using an existing structure (PDB: 1sie) after mutating residues to match the A2 strain (Stehle and Harrison, 1996). 8A7H5 Fab was initialized using a SWISS-MODEL homology model of an unrelated Fab from mouse mAb 14 (PDB: 3gk8), with manual rebuilding of the complementarity determining regions (CDRs) (Hafenstein et al., 2009; Waterhouse et al., 2018). All models were then refined using sequential rounds of manual building in COOT and automated refinement in PHENIX (Emsley et al., 2010; Liebschner et al., 2019). Models were validated with MolProbity (Chen et al., 2010).

### Data and code availability

Maps and models for the pentavalent and hexavalent capsomers are deposited at XXXX for both MuPyV (PDB XXXX, XXXX) and the MuPyV-Fab complex (PDB XXXX, XXXX). The icosahedral maps are likewise deposited as EMDB-XXXX and EMDB-XXXX. ISECC, our custom subparticle extraction program, is available at on GitLab (https://git.psu.edu/suh21/isecc).

### Statistical analysis

Statistical analyses were performed using Prism 8 software (GraphPad) using a Mann-Whitney *U* test, multiple t tests with statistical significance determined by the Holm-Sidak method, or ordinary one-way ANOVA with Tukey’s multiple comparisons test. P values of <0.05 were considered significant and all data is shown as mean, with error bars representing SD. Figures contain the data from all repeats and no data points were excluded. Statistical methods were not used to pre-determine sample sizes. The quantitation of hydrocephalus was performed in a blinded manner, no blinding was employed for other experiments. All sample sizes, numbers of repeats, and statistical tests are included in the Figure Legends. In all figures, ns = p>0.05, *p<0.05, **p<0.01, ***p<0.001, ****p<0.0001. All significant differences are labeled.

