## Supplementary material for "Antibody Escape by Polyomavirus Capsid Mutation Facilitates Neurovirulence": Key Resources Table

| Reagent or Resource | SOURCE | IDENTIFIER |
|---|---|---|
| **Antibodies, Staining Reagents, and Tetramers** | | |
| Anti-VP1 (Rat Clone 8A7H5) | Swimm et al. 2010 | N/A |
| ChromPure Rat IgG | Jackson ImmunoResearch | Cat#012-000-003 |
| Anti-VP1 (Rabbit polyclonal) | Provided by Robert Garcea | N/A |
| Anti-Vimentin (Rat Clone 280618) | R&D Systems | Cat#MAB2105 |
| Anti-GFAP (Goat polyclonal) | Abcam | Cat#ab53554 |
| Anti-Iba1 (Rabbit polyclonal) | FUJIFILM Wako | Cat#019-19741 |
| Anti-CD3 (Rabbit Clone SP7) | Abcam | Cat#ab16669 |
| Anti-CD13 (Goat polyclonal) | R&D Systems | Cat#AF2335 |
| Anti-Goat IgG AF488 (Bovine polyclonal) | Jackson ImmunoResearch | Cat#805-545-180 |
| Anti-Rat IgG AF568 (Donkey polyclonal) | Abcam | Cat#ab175475 |
| Anti-Rabbit IgG AF647 (Donkey polyclonal) | Jackson ImmunoResearch | Cat#711-605-152 |
| ProLong Gold antifade reagent with DAPI | ThermoFisher | Ref#P36931 |
| Fixable Viability Dye eFluor780 | ThermoFisher | Cat# 65-0865-14 |
| Anti-CD8α-AF700 (Clone 53-6.7) | Biolegend | Cat#100730 |
| Anti-CD44-FITC (Clone IM7) | Biolegend | Cat#103006 |
| $D^b$-LT359 Tetramer | NIH Tetramer Core | N/A |
| Anti-Rat IgG-APC (Goat polyclonal) | BD | Cat#551019 |
| Anti-Mouse IgG-HRP (Poly4053) | Biolegend | Cat#405306 |
| Anti-Mouse IgG-HRP (Goat polyclonal) | Bethyl Laboratories INC | Cat#A90-116P |
| **Bacterial and Viral Strains** | | |
| MuPyV (Strain A2) | N/A | N/A |
| BL21 (*E. coli*) | Agiltent | Cat#200133 |
| **Plasmids** | | |
| PyVP1-pGEX-4T-2 | Provided by Robert Garcea | N/A |
| H2B-GFP | Kanda et al. 1998, Addgene | Plasmid #11680 |
| **Chemicals, Peptides, and Recombinant Proteins** | | |
| Benzonase® Nuclease | Sigma | Cat#E1014 |
| Neuraminidase from *Vibrio cholerae* (Type II) | Sigma | Cat#N6514 |
| OptiPrep™ | STEMCELL Technologies | Cat#07820 |
| PFHM-II Protein-Free Hybridoma Medium | ThermoFisher | Ref#12040-077 |
| Glutathione Sepharose 4B | GE Healthcare | Cat#17075601 |
| DMEM, 1X | VWR | Cat#10-013-CV |
| TRIzol® Reagant | ThermoFisher | Ref#15596018 |
| RevertAid H Minus Reverse Transcriptase | ThermoFisher | Cat#EP0451 |
| Lipofectamine™ 2000 Transfection Reagent | ThermoFisher | Cat#11668030 |
| **Critical Commercial Assays** | | |
| TBP PrimeTime® XL qPCR Assay | IDT | Mm.PT.39a.22214839 |
| 1-Step® Ultra TMB-ELISA | ThermoFisher | Ref#34028 |
| 96 Well EIA/RIA Polystyrene High Bind Microplate | Fisher Scientific | Cat#3590 |
| PerfectCTa® SYBR® Green FastMix® | Quantabio | P/N 84069 |

| | | |
|---|---|---|
| PerfectCTa® FastMix® II ROX | Quantabio | P/N 84210 |
| PureLink® Viral RNA/DNA mini Kit | ThermoFisher | Ref#12280-050 |
| Pierce™ Fab Micro Preparation Kit | ThermoFisher | Ref#44685 |
| Nab™ Protein G Spin Columns | ThermoFisher | Ref#89953 |
| Wizard® Genomic DNA Purification Kit | Promega | Ref#A1120 |
| QuikChange II Site-Directed Mutagenesis Kit | Agilent | Cat#200523 |
| CELLine Disposable Bioreactor | Fisher Scientific | Cat#353137 |
| QIAquick PCR Purification Kit | Qiagen | Cat#28104 |
| **Experimental Models: Cell Lines** | | |
| BALB/3T3 Clone A31 | ATCC | CCL-163 |
| NMuMG | ATCC | CRL-1636 |
| mIMCD-3 | ATCC | CRL-2123 |
| **Experimental Models: Organisms/Strains** | | |
| C57BL/6 Mice | National Cancer Institute | Cat#OIC55 |
| STAT1$^{-/-}$ Mice | Jackson Laboratory | Cat#012606 |
| µMT Mice | Jackson Laboratory | Cat#002288 |
| **Oligonucleotides** | | |
| V296F Mutagenesis Forward 5’-CTCTCCAGTGATGGAAATCATAGTTTCTTG-3’ | Invitrogen | N/A |
| V296F Mutagenesis Reverse 5’-CAAGAAACTATGATTTCCATCACTGGAGAG-3’ | Invitrogen | N/A |
| V296A Mutagenesis Reverse 5’-GCCCTCTCCAGTGATGGGCATCATAGTTTCTTGTAACTC-3’ | IDT | N/A |
| V296A Mutagenesis Reverse 5’-GAGTTACAAGAAACTATGATGCCCATCACTGGAGAGGGC-3’ | IDT | N/A |
| V296Y Mutagenesis Forward 5’-GGGAAGCCCTCTCCAGTGATGGTAATCATAGTTTCTTGTAACTC-3’ | IDT | N/A |
| V296Y Mutagenesis Reverse 5’-GAGTTACAAGAAACTATGATTACCATCACTGGAGAGGGCTTCCC-3’ | IDT | N/A |
| N293F Mutagenesis Forward 5’-GTGATGGACATCATAGAATCTTGTAACTCTCCAGCCCATTATATC-3’ | IDT | N/A |
| N293F Mutagenesis Reverse 5’-GATATAATGGGCTGGAGAGTTACAAGATTCTATGATGTCCATCAC-3’ | IDT | N/A |
| N293Y Mutagenesis Forward 5’- | IDT | N/A |

| | | |
|---|---|---|
| GTGATGGACATCATAGTATCTTGTAACTCTCCAGCCCATTATATC-3’ | | |
| N293Y Mutagenesis Reverse<br>5’-GATATAATGGGCTGGAGAGTTACAAGATACTATGATGTCCATCAC-3’ | IDT | N/A |
| T291D Mutagenesis Forward<br>5’-GTGATGGACATCATAGTTTCTGTCAACTCTCCAGCCCATTATATC-3’ | IDT | N/A |
| T291D Mutagenesis Reverse<br>5’-GATATAATGGGCTGGAGAGTTGACAGAAACTATGATGTCCATCAC-3’ | IDT | N/A |
| H139R Mutagenesis Forward<br>5’-CTGTGGGTTTGTTGAACCCACGCACATCTAACAGTGAGCCAGAGC-3’ | IDT | N/A |
| H139R Mutagenesis Reverse<br>5’-GCTCTGGCTCACTGTTAGATGTGCGTGGGTTCAACAAACCCACAG-3’ | IDT | N/A |
| R77N Mutagenesis Forward<br>5’-GTAGCCAAATTAATCCCATTGCTCCAACCATAGTATTGCCCTCCC-3’ | IDT | N/A |
| R77N Mutagenesis Reverse<br>5’-GGGAGGGCAATACTATGGTTGGAGCAATGGGATTAATTTGGCTAC-3’ | IDT | N/A |
| N80K Mutagenesis Forward<br>5’-CTATGGTTGGAGCAGAGGGATTAAGTTGGCTACATCAGATACAGAGGATTCCCC-3’ | IDT | N/A |
| N80K Mutagenesis Reverse<br>5’-GGGGAATCCTCTGTATCTGATGTAGCCAACTTAATCCCTCTGCTCCAACCATAG-3’ | IDT | N/A |
| Δ294 Mutagenesis Forward<br>5’-CTGGGAAGCCCTCTCCAGTGATGGACATCATTTCTTGTAACTCTCCAGCCCATTATATC-3’ | IDT | N/A |
| Δ294 Mutagenesis Reverse<br>5’-GATATAATGGGCTGGAGAGTTACAAGAAATGATGTCCATCACTGGAGAGGGCTTCCCAG-3’ | IDT | N/A |
| LT DNA qPCR Forward | IDT | N/A |

| | | |
|---|---|---|
| 5’-CGCACATACTGCTGGAAGAAGA-3’ | | |
| LT DNA qPCR Reverse<br>5’-TCTTGGTCGCTTTCTGGATACAG-3’ | IDT | N/A |
| LT DNA qPCR probe<br>5’-ATCCTTGTGTTGCTGAGCCCGATG-3’ | IDT | N/A |
| V296F DNA qPCR Detection Forward<br>5’-CCCTCTCCAGTGATGGAA-3’ | IDT | N/A |
| V296F DNA qPCR Detection Reverse<br>5’-AGAACACAAGGTACTTTGGC-3’ | IDT | N/A |
| LT mRNA qPCR Forward<br>5’-AGGAATTGAACAGTCTCTGGG-3’ | IDT | N/A |
| LT mRNA qPCR Reverse<br>5’-GTCATCGTGTAGTGGACTGTG-3’ | IDT | N/A |
| LT mRNA qPCR probe<br>5’-AACCGGCTTCCAGGGCTCT-3’ | IDT | N/A |
| VP1 Amplification and Sequencing Forward<br>5’- CGACCCCTTGAAGGACATATGTGAA-3’ | IDT | N/A |
| VP1 Amplification and Sequencing Reverse<br>5’-CACCTACTTGGGCAACAGTCA-3’ | IDT | N/A |
| **Software and Algorithms** | | |
| Prism | Graphpad | v 8.4.2.679 |
| FlowJo | BD | v 10.6.1 |
| ImageJ | NIH | v 1.8.0 |
| Leica LAS X | Leica | v 3.7.2 |
| Photoshop | Adobe | v 19.1.6 |
| Relion | Scheres et al. 2019 | v 3.0, 3.1 |
| cryoSPARC | Structura Biotechnology | v 2.8.3 |
| ISECC | See code availability | v 2019.09 |
| PHENIX | phenix-online.org | V 1.17.1-3660 |
| Coot | Emsley et al. 2010 | v 0.8.9 |
