## Supplemental Figures and Tables for "Antibody Escape by Polyomavirus Capsid Mutation Facilitates Neurovirulence"

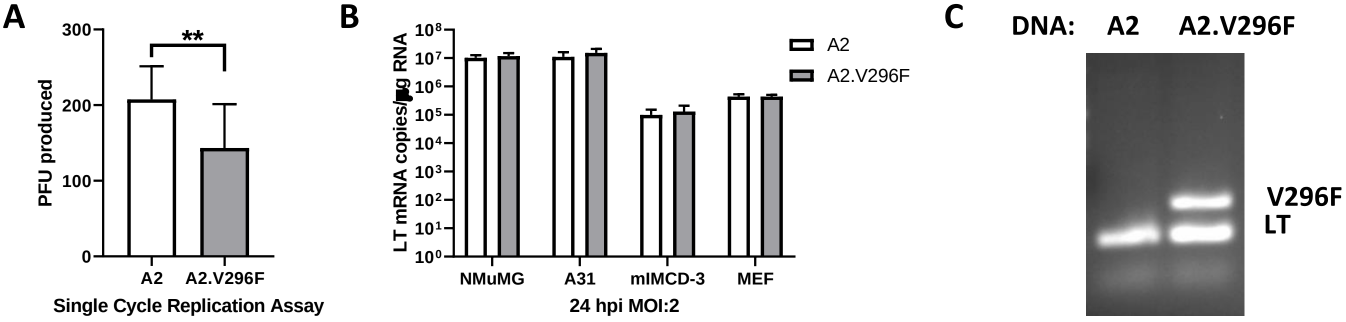

**Supplemental Figure 1. The A2.V296F VP1 mutant virus retains infectivity *in vitro*.**
(A) Single cycle replication assay in A31 fibroblasts. Cells were infected with an MOI of 0.1 and virus was collected 60 hpi and measured by plaque assay. Data are from two independent experiments, n = 15.(B) LT mRNA levels in cell lines and primary MEFs 24 hpi with an MOI of 2. Data are from two independent experiments, n = 6. (C) PCR products after amplification of A2 or A2.V296F DNA with a combination of PCR primers for LT or the V296F-VP1 mutation. Data analyzed by Mann-Whitney *U* test (A) or multiple t tests (B). $^{**}p<0.01$.

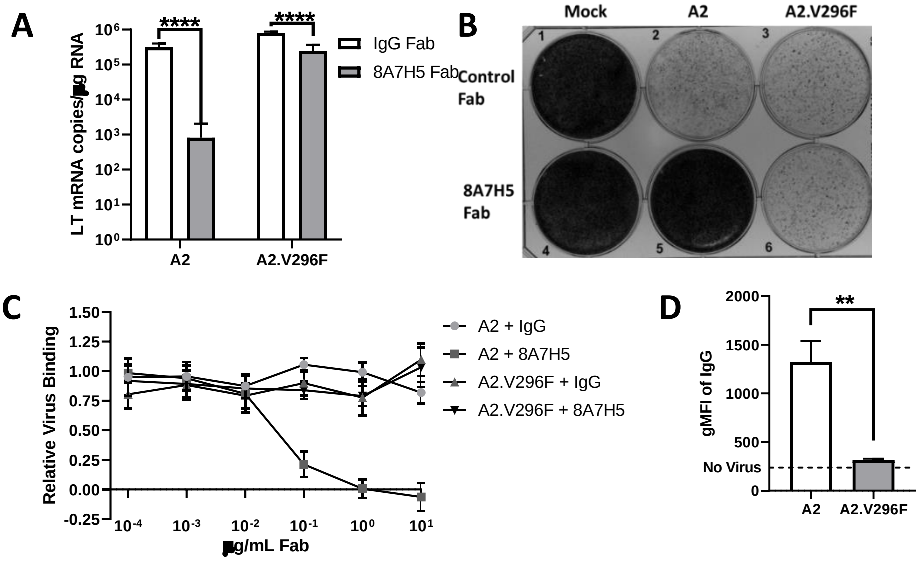

**Supplemental Figure 2. 8A7H5 Fab neutralizes A2 but not A2.V296F.**
(A) LT mRNA levels in NMuMG cells 24 hpi with A2 or A2.V296F preincubated with 8A7H5 Fab or control Fab. Data are from two independent experiments, n = 6. (B) A31 fibroblasts were infected with A2 or A2.V296F at 0.01 MOI, with 8A7H5 or control Fab added 24 hpi. Cells were fixed with formaldehyde and stained with crystal violet 7 dpi. (C) Increasing concentrations of 8A7H5 Fab prevent the attachment of A2, but not A2.V296F, to NMuMG cells. Data from two independent experiments, n = 5-6. (D) Binding of 8A7H5 Fab to VP1 is prevented by V296F. NMuMG cells were incubated with A2 or A2.V296F followed by incubation with 8A7H5 Fab. Bound 8A7H5 was detected with an anti-IgG secondary. Data are from two independent experiments, n = 6. Data analyzed by multiple t tests (A) or Mann-Whitney *U* test (B). **p<0.01, ****p<0.0001.

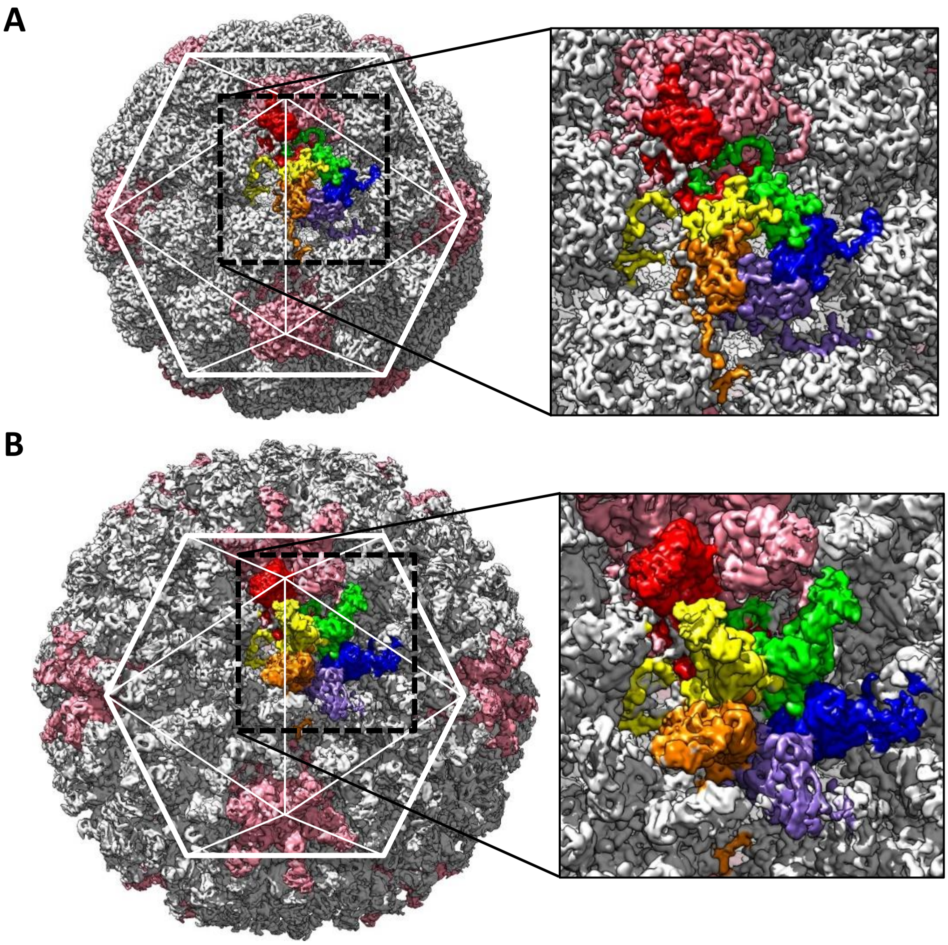

**Supplemental Figure 3. Asymmetric Unit.**
(A) The surface rendered MuPyV capsid map overlaid with an asymmetric unit (ROYGBV) consisting of one VP1 chain from the pentavalent capsomer (red), together with five VP1 chains from a neighboring hexavalent capsomer (grey). (B) The MuPyV-Fab complex map with Fab and VP1 colored as in (A). Each asymmetric unit contains six nearly identical copies of the Fab epitope.

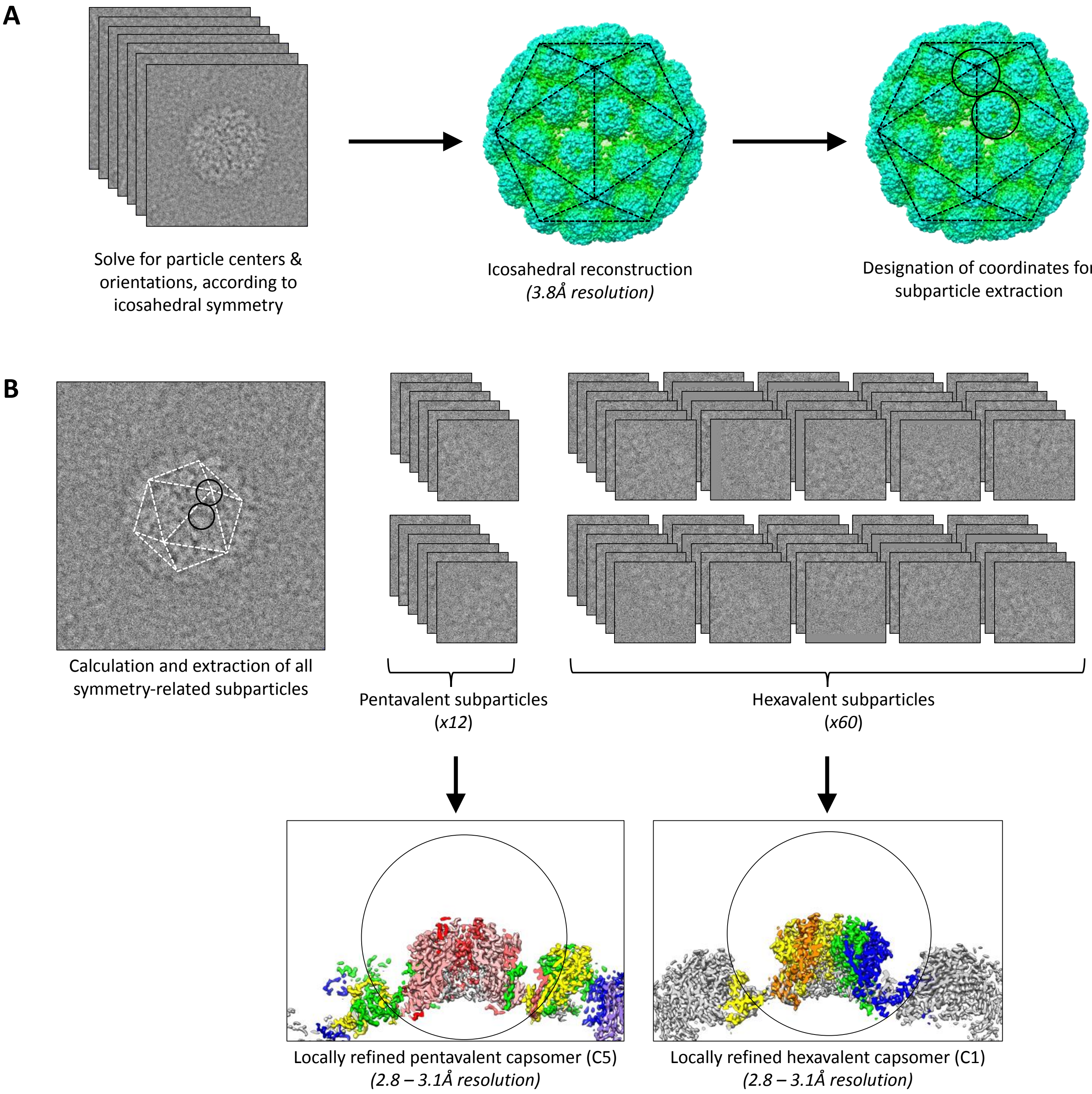

**Supplemental Figure 4. Refinement workflow.**
(A) Icosahedral refinement produces a moderate resolution map of the full capsid and allows for designation of x,y,z coordinates of pentavalent (red) and hexavalent capsomers (blue). (B) Coordinates and orientations for all pentavalent and hexavalent capsomers are calculated, which enables subparticle extraction and local refinement of capsomer centers and orientations, significantly improving map resolution compared to the initial icosahedral reconstruction.

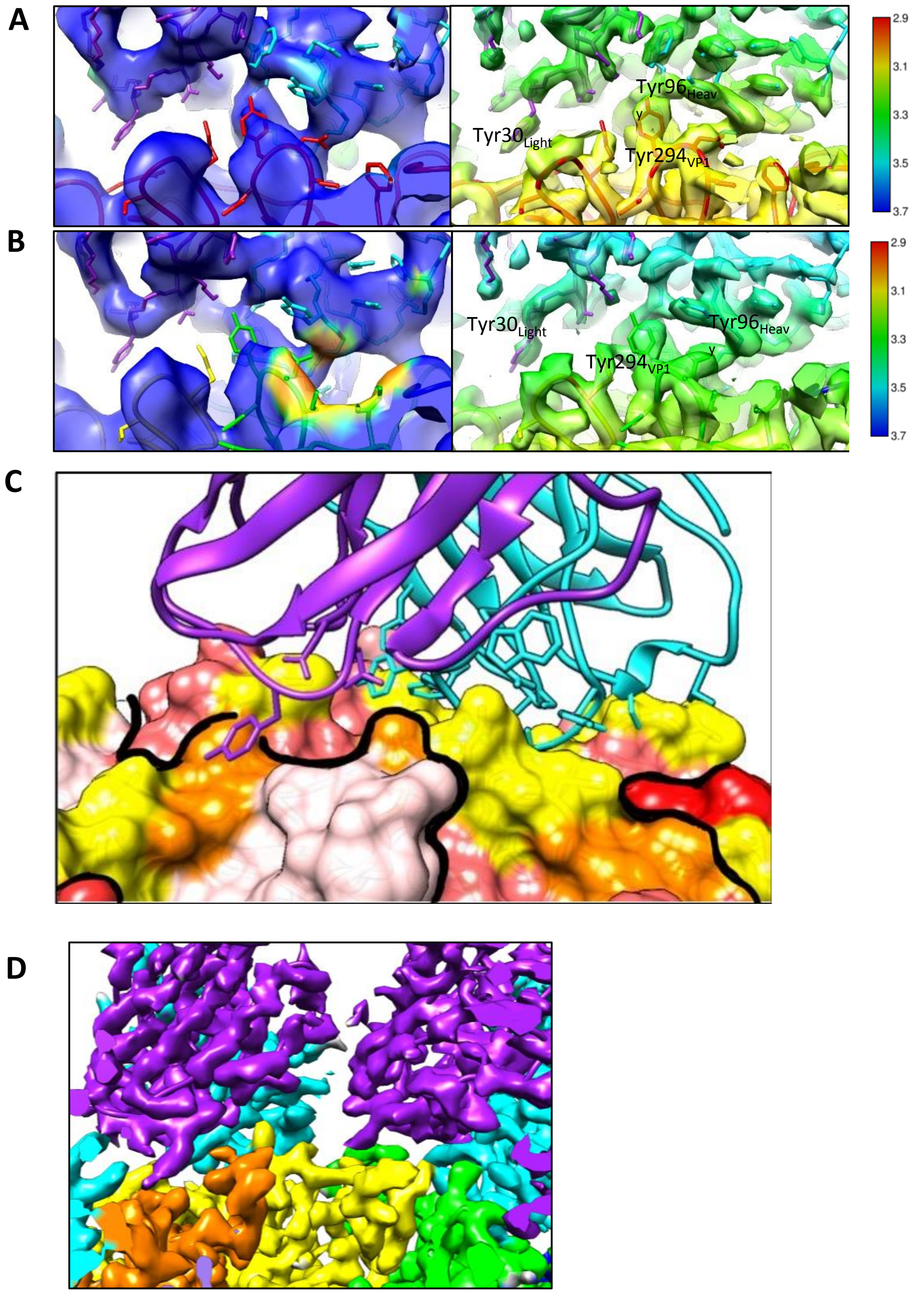

**Supplemental Figure 5. Local resolution**

(A & B) Equivalent zoomed views with density colored according to local resolution at the interface for the pentavalent (A) and hexavalent (B) capsomers show resolution improved after subvolume refinement (right) compared to the initial icosahedral refinement (left). (C) The heavy chain (cyan) interacts with one copy of VP1 (yellow residues) with the light chain (purple) making additional contacts with the neighboring VP1 (orange residues). Symmetry-related VP1 molecules are shown in shades of red, divided by a thick black line. (D) Variable domains of adjacent bound Fab do not clash (heavy: cyan; light: purple). VP1 chains are colored as in Fig 4 D,E.

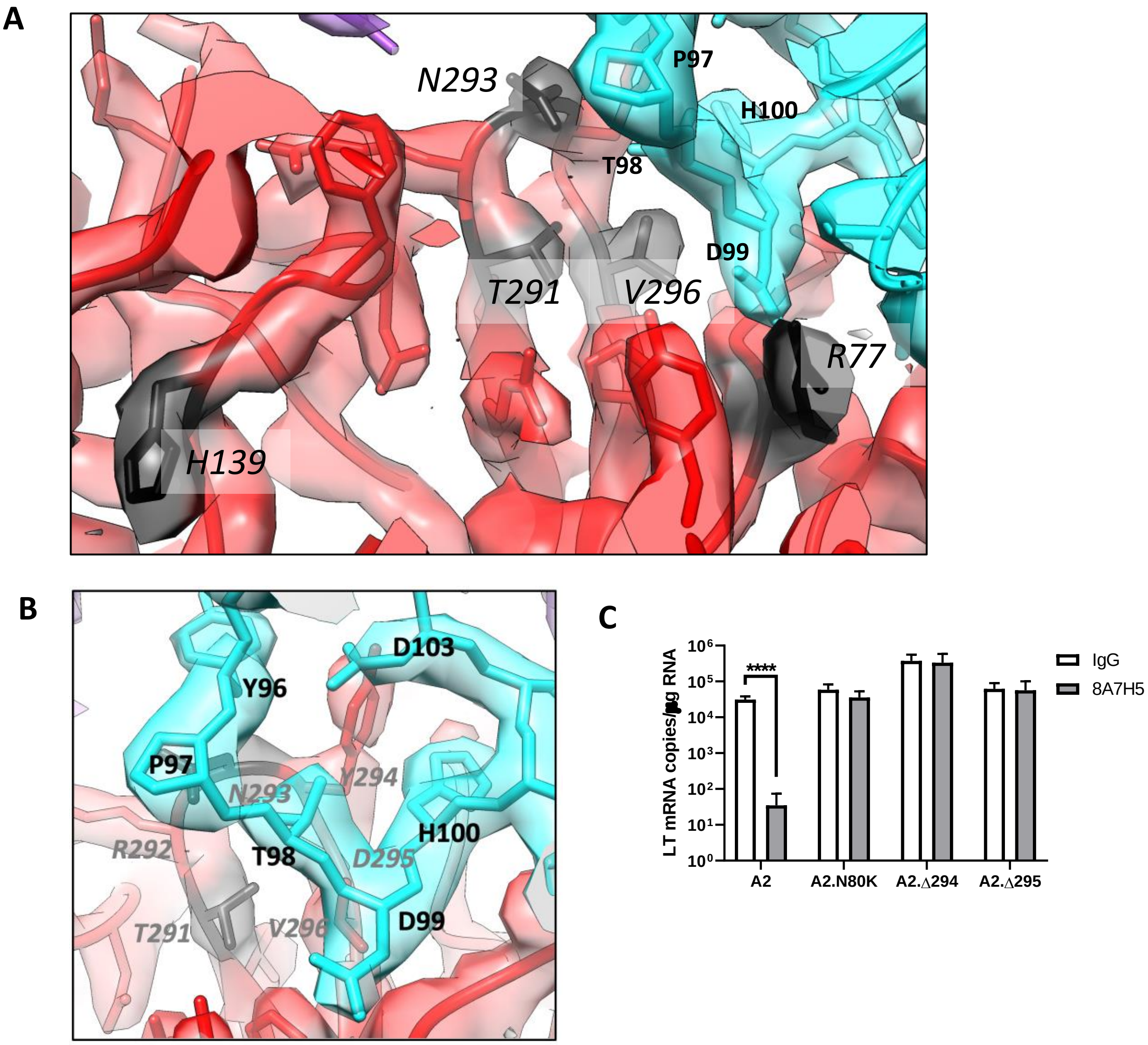

**Supplemental Figure 6. Relation of other VP1 mutations to virus-Fab interface.** (A) Cryo EM density at the interface between the pentavalent capsomer (red) and Fab (cyan, purple). Location of the residues mutated to mimic those found in JCPyV-PML (H139, T291, N293, V296, R77) are noted in black. Residues T291, N293, V296 and R77 are located at the interface whereas H139 is a buried residue. (B) An alternate view of the interface shows the interaction between the HI loop and the Fab heavy chain CDR loop H3. (C) LT mRNA levels in NMuMG cells 24 hpi with A2, A2.N80K, A2.Δ294, A2.Δ295 preincubated with 8A7H5 or control IgG. Data are from two independent experiments, n = 6. Data were analyzed by multiple t tests (C). ****p<0.0001.

| | MuPyV | MuPyV-Fab |
|---|---|---|
| Microscope/Detector | Titan Krios | |
| Detector | Falcon 3 | |
| Magnification | x59,000 | |
| Voltage (kV) | 300 | |
| Electron exposure | 45 | |
| Defocus range (µm) | 1. – 3.0 |  |
| Pixel Size | 1.1 Å | |
| Micrographs (total) | 1,756 | 1,811 |
| Micrographs (used) | 1,750 | 1,804 |
| Particles (total) | 17,416 | 9,811 |
| Particles (used) | 15,499 | 9,146 |

**Supplemental Table 1. Collection statistics.** Independent datasets were collected for MuPyV and the MuPyV-Fab complex.

| Sample | MuPyV | | | MuPyV-Fab | | |
|---|---|---|---|---|---|---|
| Map composition | capsid | capsomer (pentavalent) | capsomer (hexavalent) | capsid | capsomer (pentavalent) | capsomer (hexavalent) |
| Particle number | 15,499 | 185,988 | 929,940 | 9,146 | 109,752 | 548,769 |
| Symmetry imposed | I1 | C5 | C1 | I1 | C5 | C1 |
| Map resolution (Å) | 3.9Å | 2.9 | 2.9 | 4.2Å | 3.1 | 3.2 |
| Local resolution (Å) | | 2.7 – 3.1 | 2.7 – 3.1 | | 2.9 – 3.8 | 3.0 – 3.8 |
| Model composition | | VP1 | VP1 | | VP1, 8A7H5 Fab | VP1, 8A7H5 Fab |
| VP1 | | chain F (x5) | chain A-E | | chain F (x5) | chain A-E |
| Protein residues | | 1660 | 1687 | | 1660 | 1687 |
| RMS bonds (Å) | | 0.0044 | 0.0043 | | 0.0073 | 0.0078 |
| RMS angles (°) | | 0.76 | 0.72 | | 0.82 | 0.84 |
| Validation | | | | | | |
| Molprobity Score | | 2.97 | 2.67 | | 2.86 | 2.82 |
| Clashscore | | 11.99 | 9.16 | | 11.59 | 9.98 |
| Rotamer outliers (%) | | 11.60 | 8.09 | | 9.29 | 9.10 |
| Ramachandran plot | | | | | | |
| Favored (%) | | 89.43 | 91.91 | | 90.06 | 89.33 |
| Outliers (%) | | 0.29 | 0.39 | | 0.29 | 0.34 |
| Fab | | | | | 1x | 5x |
| Protein residues | | | | | 215 | 1070 |
| RMS bonds (Å) | | | | | 0.0043 | 0.0048 |
| RMS angles (°) | | | | | 0.74 | 0.85 |
| Validation | | | | | | |
| Molprobity Score | | | | | 2.95 | 3.35 |
| Clashscore | | | | | 18.74 | 16.71 |
| Rotamer outliers (%) | | | | | 8.33 | 19.27 |
| Ramachandran plot | | | | | | |
| Favored (%) | | | | | 92.31 | 85.78 |
| Outliers (%) | | | | | 0.00 | 0.46 |

**Supplemental Table 2. Refinement statistics.** Local refinement allowed models to be built into the higher resolution capsomer maps.

| CDR | | | **VP1 residue** |
|---|---|---|---|
| L1 | Tyr | 30 | 141, 292 |
| L3 | Asn | 92 | 83 |
| | Ala | 93 | 83 |
| H1 | Ser | 26 | 67, 68 |
| H2 | Ser | 49 | 77 |
| | Ala | 50 | 77 |
| | Asp | 52 | 91 |
| | Asn | 69 | 68 |
| H3 | Tyr | 96 | 293 |
| | Thr | 98 | 293 |
| | Asp | 99 | 77, 78, 80 |
| | His | 100 | 80, 151, 294, 296 |
| | Phe | 101 | 80 |
| | Tyr | 102 | 83 |
| | Asp | 103 | 294 |
| | Trp | 104 | 294 |

**Supplemental Table 3. Fab contact residues.** The majority of Fab contacts are through the heavy chain, with minor contributions from the light chain.

| **Virus** | Avian polyomavirus | BK polyomavirus | BK polyomavirus | BK polyomavirus | Murine polyomavirus | Murine polyomavirus |
|---|---|---|---|---|---|---|
| **Author** | Shen et al. | Hurdiss et al. | Lindner et al. | Hurdiss et al. | This study | This study |
| **PMID** | 21239031 | 26996963 | 30824324 | 29706532 | -- | -- |
| **PDB** | 3IYS | 5FUA | 6GG0 | 6ESB | -- | -- |
| **Year** | 2011 | 2016 | 2019 | 2018 | 2020 | 2020 |
| **Ligand** | N/A | N/A | scFv | oligosacch. | Fab | none |
| **Resolution** | 11.3Å | 7.6Å | 4.2Å | 3.4Å | 3.1Å [2.9 – 3.8Å] | 2.9Å [2.7 – 3.1Å] |
| **Particles** | 5,338 | 2,237 | 5,000 | 40,334 | 9,171 | 15,499 |
| **Pixel Size** | ?? | 1.9Å | 1.1Å | 1.1Å | 1.1Å | 1.1Å |
| **Microscope** | Tecnai-F30 | Tecnai-F20 | Krios | Krios | Krios | Krios |
| **Detector** | film | K2 | K2 | Falcon III | Falcon III | Falcon III |

**Supplemental Table 4. Statistics of existing and new cryo EM polyomavirus maps.** Capsomer-based local refinement allowed rapid refinement of polyomavirus to high resolution, using only a modest particle number.
